# Intrinsic disorder in CENP-A^Cse4^ tail and its chaperone facilitates synergistic association for kinetochore stabilization

**DOI:** 10.1101/2022.08.15.504061

**Authors:** Shivangi Shukla, Anusri Bhattacharya, Prakhar Agarwal, Nikita Malik, Kalaiyarasi Duraisamy, Nithyakalyani Sri Rangan, Ramakrishna V. Hosur, Ashutosh Kumar

## Abstract

The kinetochore is a complex multiprotein network that assembles at a specialized DNA locus called the centromere to ensure faithful chromosome segregation. The centromere is epigenetically marked by a histone H3 variant – the CenH3. The budding yeast CenH3, called Cse4, consists of an unusually long and disordered N-terminal tail that has a role in kinetochore assembly. Its disordered chaperone, Scm3 is involved in its centromeric deposition as well as in the maintenance of a segregation-competent kinetochore. The dynamics of the Cse4 N-tail and chaperone interaction have not been studied, leaving a gap in our understanding of their roles at the centromere. Previously, we had shown that Scm3 is an intrinsically disordered protein. Here, using NMR and a variety of biophysical and bioinformatics tools, we show that Cse4 N-tail is also disordered, the two proteins interact with each other at multiple sites, and this interaction reduces the disorder in Scm3; the chain opens up relative to the native state ensemble and the conformational exchange is reduced. Interestingly, this interaction between the two intrinsically disordered protein is fairly specific as seen by positive and negative controls, and is majorly driven by electrostatics as both the proteins have multiple acidic and basic regions. The complex retains a fair amount of disorder, which facilitates a synergistic association with the essential inner kinetochore Ctf19-Mcm21-Okp1-Ame1 complex; a model has been suggested to this effect. Given the abundance of intrinsic disorder in the kinetochore proteins, this type of interaction and adaptation may be prevalent in other proteins as well for mediating kinetochore assembly. Thus, the present study, on one hand, provides significant structural and mechanistic insights into the complex and dynamic process of kinetochore assembly, and on the other hand, illustrates a mechanism that intrinsically disordered proteins would adapt to mediate the formation of complex multiprotein networks, in general.

**Graphical Abstract:** 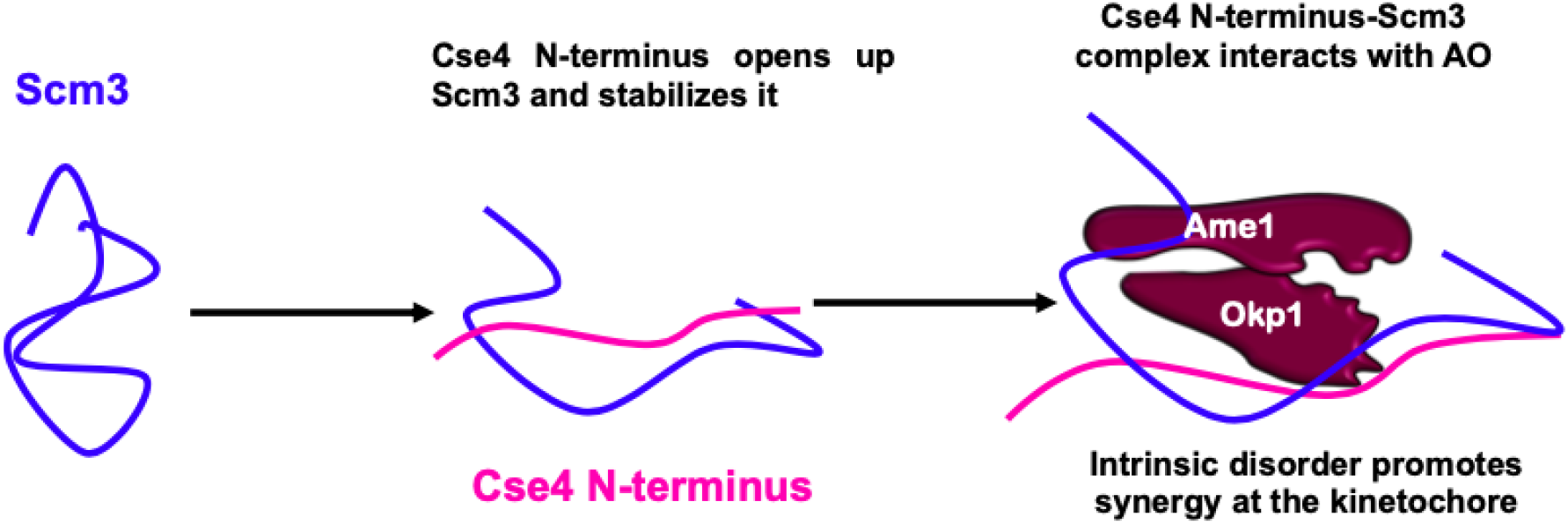

## Introduction

Faithful segregation of chromosomes is a pre-requisite for successful cell division. Errors in mitotic chromosome segregation lead to aneuploidy and cancer (1) whereas faults during meiosis leads to various developmental defects (2). In eukaryotes, a specific DNA locus called the centromere ensures accurate cell division. Centromeres are epigenetically marked by the presence of a specialized nucleosome wherein Histone 3 (H3) is replaced by a variant identified as CENP-A (Centromere Protein A) in humans (3), Cse4 (Chromosome Segregation 4) in *S. cerevisiae* (4), CID or CenH3 (Centromeric Histone 3) in *Drosophila* (5). The CENP-A nucleosome acts as a scaffold for the assembly of an essential multiprotein complex called the kinetochore that links chromosomes to spindle microtubules (6). The budding yeast centromere is defined by a 125 bp DNA sequence and consists of a single Cse4 containing nucleosome (7). Cse4 consists of a conserved histone-fold domain (HFD) at the C-terminus and an extended N-terminal tail (∿129 amino acids). The N-terminal tail of Cse4 is unique as it does not contain any sequence homology to the H3 N-tail (8). The Cse4 HFD is crucial for centromere localization and has been at the center of structural studies while the N-terminal tail appears to have no direct role in centromere targeting (9). An essential non-histone protein called Scm3 acts as a specific chaperone to Cse4 and targets it to the centromere by recognizing the centromere-targeting domain (CATD) in the HFD of Cse4 (10–13). Structural studies on truncated protein constructs of Scm3,Cse4 and H4 reveal that Scm3 forms extensive contacts with both Cse4 HFD and H4 and induces conformational changes that prevents DNA binding of H4 (14–16). Hence, for proper assembly of the centromeric nucleosome, Scm3 has to be evicted from the Cse4-H4-Scm3 heterotrimeric complex. Notably, the entire N-terminal tail of Cse4 was deleted in the construct, thus presenting a knowledge gap in the understanding of the interaction dynamics at the centromere. Scm3 is intrinsically disordered in solution and undergoes a disorder-to-helical transition in presence of Cse4 HFD (12). It should be noted here that in addition to being a targeting and assembly factor for Cse4 (17), Scm3 is also imperative for cell cycle progression. Scm3-depleted cells undergo a Spindle assembly checkpoint (SAC) dependent cell-cycle arrest (10) as well as reduced localization and recruitment of essential inner KT proteins such as Mif2 and Ndc10 to form and maintain a segregation-competent KT (18). Hence, the dislodged Scm3 may remain associated with the centromere via Ndc10/DNA binding and/or through interaction with other proteins such as the Cse4 N-terminus at the centromere. It has been shown that Scm3 dynamically exchanges at the centromere (19) and we speculate that this will allow Scm3 to interact with different proteins at different stages of the cell cycle. With the N-terminal tail of Cse4 freely available for interactions and Scm3 being in a dynamic exchange at the centromere, does Scm3 also associate with the N-tail for persistent centromeric association? If yes, does it perform any function in association with the N-tail?

It has been suggested previously that the N-and C-terminus of Cse4 behave as distinct domains and may have separate folding pathways and functions (20). Further, it has been demonstrated by our group that Cse4 exists in a ‘closed conformation’ via self-interaction between the N-and C-terminal domains. Upon interaction with H4, the N-terminal tail is released (21) which then becomes available for PTMs as well as interaction with proteins critical for centromere functions, including kinetochore protein recruitment and assembly, kinetochore-microtubule connections, and sister chromatid cohesion. Specifically, the essential domain of the N-terminus (END, residues 28-60) has been shown to be involved in an interaction with the essential Ame1/Okp1(AO) heterodimer of the CENP-P/Q/O/U ^Ctf19p-Mcm21p-Okp1p-Ame1p^ (COMA) complex (22, 23) indicating that N-terminus has a role in KT assembly. AO at the N-terminus and Mif2 at the C-terminus of Cse4 are two binding partners that mediate KT assembly apart from CENP-T^Cnn1^ (24).

Understanding the mechanisms underlying complex protein networks such as the KT involves studying its components separately and together. Intrinsic disorder has emerged as a paradigm shift in the structure-function relationship and painting a complete picture of the KT would require the knowledge of its disordered counterparts. Because there is no definite 3D structure, **I**ntrinsically **D**isordered **P**roteins/**R**egions (IDP(R)s) span a larger area and can thus interact with proteins or other biomolecules (DNA, RNA) that are further away. They can also provide a large interaction interface for multiple binding partners while minimizing steric interference (25). This IDP(R) property is used to mediate multiprotein complex assemblies. Additionally, they can also sample a large conformational space and quickly interconvert between states, resulting in significant heterogeneity. IDP(R)s gain promiscuity as a result of heterogeneity, allowing them to bind to multiple ligands, making them ideal candidates for mediating multiprotein interactions (26).

## Materials and methods

### Cloning of Cse4 N-terminus protein

Cse4 N-terminus (1-129) was cloned in pGEX6p1 vector with ampicillin resistance. The sequences of primers used for the cloning were 5’-GGATCCCATATGTCAAGTAAACAACAATGGGTTAG-3 (forward) containing BamHI and NdeI restriction sites and 5’-CTCGAGTTATTCGACGCGCTTTAAGCTCTGC-3’ (reverse) harboring the XhoI restriction site. The clones contained a GST tag with a PreScission cut site. The clone was verified by sequencing (Eurofins Scientific) and used for further studies.

### Purification of GST-tagged Cse4 N-terminus

The N-terminus plasmid was transformed into E. *coli* BL21 DE3 cells for protein expression. A single colony of the recombinant bacteria was inoculated in 10 ml LB medium supplemented with ampicillin and was incubated overnight at 37°C shaker incubator. 10ml of the culture was transferred into 1L LB medium supplemented 100µg/mL Ampicillin & incubated at 37°C in a shaker incubator. 0.5 mM IPTG was added to the culture as the O.D reached 0.6 and was then incubated for 6 hrs following which it was centrifuged at 9000g for 10 min. The cell pellet was resuspended in Lysis buffer (20mM sodium phosphate pH8, 100mM NaCl, 1mM EDTA, 1% Triton X-100 pH 8). The resuspended cells were sonicated for 15mins and centrifuged at 20000g for 40 mins to obtain a clear supernatant containing the recombinant protein. The supernatant was applied onto a pre-equilibrated column containing Protino Glutathione Sepharose™ 4B beads (Takara) and kept for binding for 3 hrs at 4°C. After binding, the flow through was collected and the beads were washed with wash buffers containing increasing amount of salt (20mM sodium phosphate pH8, 1mM EDTA and100mM, 300mM, 500mM) to wash off non-specifically bound proteins and impurities. The beads were again equilibrated with wash buffer containing 100mM NaCl and PreScission enzyme was added to cleave off the GST tag from the protein. The beads were incubated overnight with the enzyme at 4°C. Next day, the protein was eluted in wash buffer with 100 mM salt and the purity was checked by SDS-PAGE. Second round of purification was done in cases where the first flow-through still contained unbound GST tagged protein. The pure protein obtained was concentrated using a 3 kDa Amicon® Ultra Centrifugal Filter (Merck).

### Purification of His-tagged Scm3

The plasmid for Scm3 was a kind gift from Prof. Luger (University of Colorado Boulder). The protein was purified as described previously (27). Briefly, E. *coli* BL21 DE3 cells transformed with Scm3 plasmid were lysed by sonication and centrifuged at 20000g for 40 min. The supernatant was applied to Ni-NTA beads (Qiagen Ni-NTA agarose beads). The beads were washed with TN buffer (50mM Tris HCl pH 8 and 150mM NaCl) at room temperature and then incubated at 4°C. All further washes were done at 4°C with increasing concentrations of imidazole (25mM and 50 mM) before eluting the protein with 250 mM imidazole and the purity was checked by SDS-PAGE. The protein was dialyzed in 20 mM phosphate pH 7.4 and 500 mM NaCl to remove imidazole and concentrated using a 10 kDa Amicon® Ultra Centrifugal Filter (Merck).

### Purification of inner kinetochore protein Ame1/Okp1 dimer

The plasmid for AO was a kind gift from Prof. Westermann (University of Duisburg-Essen). The protein was purified as described previously (28). Briefly, E. *coli* BL21 RIPL cells transformed with the AO plasmid was centrifuged at 8,000 rpm for 10 min and the cell pellet was resuspended in Lysis buffer (20mM HEPES pH 8, 300mM NaCl, 30 mM Imidazole). The resuspended cells were sonicated for 15mins and centrifuged at 16000rpm for 40 mins to obtain clear supernatant containing the protein. The supernatant was applied to Ni-NTA beads (Qiagen Ni-NTA agarose beads). The beads were washed with 150 ml of wash buffer 1 (50mM HEPES, 300 mM NaCl and 30 mM Imidazole) at 4°C. The beads were extensively washed with 50 mM concentration of imidazole before eluting the protein with 300 mM imidazole. The protein was dialyzed in 20 mM phosphate pH 7.4 and 500 mM NaCl to remove imidazole. The protein was then concentrated and subjected to gel filtration using a Superdex 200 10/300 column. Pure fractions were collected and pooled together for studies.

### Purification of Histones

The plasmid for histones H2A, H2B, H3 and Cse4 was a kind gift from Prof. Luger (University of Colorado Boulder). The protein was purified as described previously (29). Briefly, *E*.*coli* BL21 RIPL cells transformed with histone plasmids were sonicated in wash buffer containing 500 mM Tris-Cl, 100 mM NaCl, 1mM EDTA and 1mM PMSF for lysis. The pellet fraction (inclusion bodies) obtained after cell lysis has the protein. The inclusion body containing histones, was washed with wash buffer + 1% Triton-X thrice to remove impurities followed by two washes of wash buffer without Triton-X. The washed pellet was then solubilized in denaturing buffer pH 7.4 overnight. The supernatant obtained after centrifugation at 20000g, 20 mins,16°C was loaded on a S200 gel filtration column to obtain pure histone fractions. Ion exchange chromatography was performed where necessary to obtain highly purified histone fractions. For refolding, DNA-free pure fractions were pooled together and dialyzed against PSB (20 mM phosphate buffer, 500 mM NaCl, and 15 mM β-ME,0.5 M urea). CD spectra were recorded after dialysis to confirm proper protein refolding.

### Purification of α-synuclein

The protein was purified as described previously (30). Briefly, *E*.*coli* BL21 cells transformed with α-synuclein plasmid were harvested by centrifugation (at 9000g rpm for 10 min at 4°C) and then were resuspended in lysis buffer containing Triton-X, lysozyme and PMSF. The cell lysate obtained was subjected to sonication for cell lysis followed by incubation at 95°C water bath for 20 mins. After cooling the cell lysate at RT, it was centrifuged at 20000g for 40 mins at 4°C. The supernatant was transferred in a fresh tube and Streptomycin sulphate (136 µl/ml) and Glacial acetic acid (228 µl/mL) were added to it. This mixture was incubated at 4°C for 30 mins and then centrifuged at 20000g for 30 mins at 4°C. To this supernatant, equal volumes of saturated ammonium sulphate was added. This mixture was then incubated at 4°C on a stirrer overnight. The next day, this solution was centrifuged at 20000g rpm for 30 mins at 4°C. The pellet obtained was resuspended in 4mL ammonium acetate and stored at 4°C. The LMW form of α-Syn was isolated using Amicon Ultra 100 kDa cutoff filters (Millipore) according to the previously described method.

### Circular dichroism spectroscopy

20 μM of the protein samples (Cse4 N-terminus, Scm3, H3 and AO) or 1:1 protein complexes of N-terminus-Scm3, H3-Scm3 and AO-Scm3 were placed in a quartz cuvette (Starna, Hainault, London) of 0.1cm path-length and the spectra were acquired in the wavelength range of 202-260 nm on a JASCO-810 instrument. Each sample was scanned thrice (3 accumulations) and the average was considered. All measurements were carried out at 25°C. The raw data was processed by smoothening and blank subtraction and the CD spectra were plotted. Three independent measurements were acquired for each sample. For complexation, the proteins were incubated in equimolar concentrations (10 μM) in 20 mM sodium phosphate, 500 mM NaCl, 5 mM DTT, pH 7.4 buffer at 25°C for 20 minutes before acquiring the spectra.

### Tryptophan fluorescence emission

Fluorescence experiments were recorded on a Fluoromax-4 spectrofluorometer equipped with a data recorder and a temperature-controller. 20 μM of Cse4 N-terminus and Scm3 in buffer 1 were taken in a 1 cm path length cuvette and excited at 295 nm and the fluorescence emission was recorded in the wavelength range of 305-400 nm at 25°C. The excitation and emission slit widths were kept at 3 nm.

### Pull-down assay

The direct interaction between Cse4 N-terminus and its chaperone Scm3 was established using a pull-down assay. Briefly, the cell pellet of Scm3 was resuspended in lysis buffer (10 mM Tris-Cl, 300 mM NaCl, 8M urea, pH 8), sonicated and centrifuged at high speed to obtain the cell supernatant. The supernatant (2mg/ml) was loaded onto a pre-equilibrated Ni-NTA column and kept for incubation for 20-30 min. The resin was then washed thoroughly with buffers (50 mM Tris-Cl, 150 mM NaCl, pH 8.0) containing increasing concentrations of imidazole (25 and 50 mM). 500µL of purified 50µM Cse4 N-terminus was then kept for binding to the column under shaking conditions for 30 min at room temperature. Subsequently, the supernatant containing the unbound Cse4 N-terminus was removed by centrifugation and the bound protein was eluted from the column using a buffer containing 250 mM imidazole. The elution products were then run on 12% SDS-PAGE to detect the fractions containing Cse4 N-terminus-Scm3 complex. Similarly, for histones H3, H2A and H2B, pull-down assay was performed with Scm3 as described above.

### Gel filtration studies

Analytical size exclusion chromatography was used to analyze the interaction between Cse4 N-terminus and Scm3. Briefly, the column containing Sephadex G-50 resin was washed and equilibrated with Buffer 1 containing 20 mM phosphate, 500 mM salt and 5mM DTT and blue dextran was passed through the column to determine the void volume. Standard proteins such as RNAase, carbonic anhydrase and ovalbumin which corresponded to the sizes of Cse4 N-terminus, Scm3 and their complex, respectively were run to estimate the fraction number of the elution peak fractions. Individually 100 µM each of Cse4 N-terminus, Scm3 and Cse4 N-terminus-Scm3 complex (1:1) were loaded onto the gel filtration column and protein fractions showing absorbance at A280 were collected and run on 12% SDS-PAGE. Graphically a plot of A280 versus elution volume was done for each of the proteins as well as the complex to depict the co-elution of the complex at the expected molecular mass.

### Fluorescent labeling of proteins

The fluorescent labeling of Cse4 N-terminus, full length Cse4, H3, Scm3 and Ame1/Okp1 complex were done using fluorescein isothiocyanate (FITC). Briefly, three-fold excess FITC dye was added to the proteins, pH 7.4 and incubated at 4°C for 4 hrs. The reaction was quenched by the addition of 1M Tris, pH 8. The excess dye was removed from the protein in a two-step process: firstly, dialyzing against phosphate buffer, pH 7.4 and then by buffer exchange using a 10kDa concentrator (Amicon, Millipore). The concentration of the labelled protein was estimated by measuring the absorbance at 495 nm using a molar extinction coefficient of 77,000 M^-1^cm^-1^ and the total protein concentration was measured at 280 nm. The incorporation ratio was determined to be ∼ 0.3.

### Determination of dissociation constant of the interaction between proteins

The dissociation constant of the interaction of Cse4 N-terminus with its chaperone Scm3 was estimated by the fluorescence-based method, as described previously (21). 100 nM FITC-Cse4 N-terminus was incubated without and with varying concentrations of Scm3 (50-5000nM) in Buffer 1 for 30 min at 25°C. The emission spectra were recorded in the range of 500-600 nm using 495 nm as the excitation wavelength. The corresponding blank spectra were also taken. The dissociation constant (kd) of the interaction between Cse4 N-terminus and Scm3 was determined by fitting the fluorescence data into the following binding isotherm:

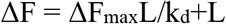

where ΔF is the change in the fluorescence intensity of FITC-Cse4 N-terminus in the presence of Scm3, ΔF_max_ is the change in the fluorescence intensity when Cse4 N-terminus is saturated with Scm3, L refers to ligand concentration i.e. concentration of Scm3. The change in the fluorescence intensity of each of the reaction set was fit into equation above using GraphPad Prism 5 (GraphPad Software). The experiment was performed three times using a Fluoromax-4 spectrofluorometer and the average kd of the three data sets was reported with the standard deviation.

### Fluorescence-based binding assays

A 1:1 nanomolar complex of FITC-labelled AO and unlabelled Cse4 N-terminus was made by incubating both the proteins at RT for 30 mins as previously described (22). The complex was titrated with increasing concentrations of Scm3 and the fluorescence intensity was recorded in triplicates. A mean and standard deviation of the three readings was used to plot the bar graphs. The experiment was performed thrice. A t-test was conducted to determine significance between the readings. The same was repeated for FITC-Scm3+AO-N-terminus; FITC-N-terminus+AO-Scm3; FITC-N-terminus+Scm3-AO and FITC-Scm3+N-terminus-AO.

### Bioinformatics analysis

Sequence-based residue-wise charge analysis of IDP ensemble of Cse4, Scm3, H3 and lysozyme was performed using CIDER (http://pappulab.wustl.edu/CIDER/analysis/).IDP ensemble was modelled using the CS23D2.0 server (http://www.cs23d.ca/index.php) and ITASSER server (https://zhanggroup.org/I-TASSER/). Secondary structure prediction of all proteins was done using IUPRED2A workbench (https://iupred2a.elte.hu/). Kappa Z-scores were calculated based on the work of Cohan and Shinn, using a code provided at https://github.com/holehouse-lab/supportingdata/tree/master/other/misc_code/kappa_z_score.

### ANS binding of Cse4 N-terminus and Scm3

ANS (8-anilinonaphthalene-1-sulfonic acid) fluorescence intensity measurements have been widely used for measuring changes in hydrophobicity in protein samples (Semisotnov GV et. al., 1991). Briefly, 500 nM of Cse4 N-terminus without and with varying concentrations of Scm3 (200,400 and 800 nM) and 800 nM Scm3 only were mixed with 20 µM ANS reagent. The samples were further incubated at RT for 30 min in dark. Post-incubation, the samples were excited at 370 nm and emission spectra were recorded between 400 and 600 nm using a Fluoromax-4 spectrophotometer. The ANS fluorescence intensities at 520 nm were plotted against incubation time.

### *CEN3* DNA preparation

Standard PCR conditions were used to prepare the centromere (*CEN3*) DNA sequence. Whole genomic DNA was isolated from wild-type yeast cells and was used as a template. Primers specific to *CEN3* sequence (GM107, GM108) were used for amplification. Following PCR, the presence of the amplicon was confirmed by agarose gel electrophoresis. The primer sequences used for the amplification are listed below:

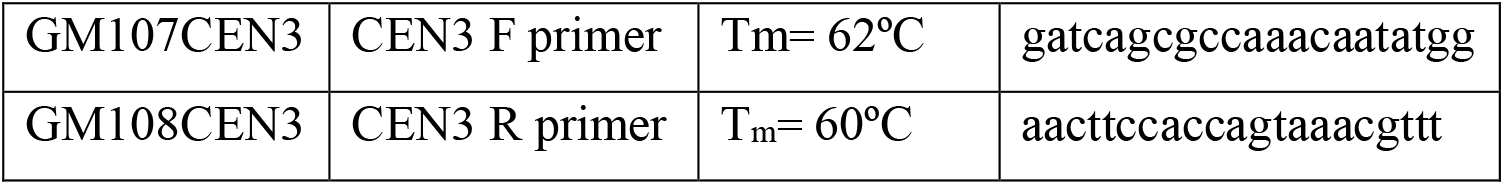

### NMR spectroscopy

#### Backbone assignment of Cse4 N-terminus

For resonance assignment, Cse4 N-terminus was produced by growing *E*.*coli* in M9 minimal media with ^15^NH4Cl and ^13^C_6_ glucose as the sole nitrogen and carbon source respectively for ^15^N and ^13^C labeling. The backbone assignment of Cse4 N-terminus was done by recording a series of 2D and 3D NMR experiments. Uniformly labeled ^13^C/^15^N Cse4 N-terminus was mixed with D2O in the ratio of 90:10 (H2O/D20) before recording the multi-dimensional NMR experiments. The experiments were recorded at 298 K on Bruker 750 MHz spectrometer (Bruker Biospin, Switzerland) using 5 mm TXI probe equipped with Z-gradient and deuterium decoupling. 2D experiments include ^15^N-^1^H HSQC and ^15^N-edited TOCSY-HSQC and the 3D experiments recorded were HNCACB, HN(CO)CACB, HNCO, HN(CA) CO, HNN (with 30% NUS), CC(CO)NH and HCC(CO)NH using standard pulse programs. The processing of the spectra were done using Topspin 3.2 and analyzed using SPARKY. The referencing for ^1^H chemical shifts was done using 2, 2-dimethyl-2-silapentane-5-sulphonic acid (DSS) and the ^13^C and ^15^N chemical shifts were referenced according to BMRB protocol.

#### Prediction of the secondary structure of Cse4 N-terminus

The secondary structure of Cse4 N-terminus was determined from the cumulative (Cα, Cβ and CO) secondary chemical shift values for each of the assigned residues (31). Sequence corrected random coil chemical shifts were used for the calculation and the secondary structure propensities were calculated according to the equation: Δδ(Ccum) = Δ(Cα)/25 − Δδ(Cβ)/25 + Δδ(CO)⁄10 where Δδ(Cα), Δδ(Cβ) and Δδ(CO) are the deviations in the chemical shifts of Cse4 N-terminus from their corresponding random coil values (32, 33).

#### NMR titration studies

The residue specific interaction regions of Scm3 with Cse4 N-terminus were identified by NMR experiments. All NMR experiments were performed on Bruker Ascend 750 MHz spectrometer at 288K with 5 mm TXI probe. Briefly, ^15^N-^1^H HSQC spectra of Scm3 in Buffer 1 was recorded in absence and presence of 1:2 equivalent of Cse4 N-terminus. The extent of interaction between the two interacting proteins was analyzed by Chemical Shift Perturbation (CSP). Perturbation of amide cross-peaks chemical shifts during the interaction studies with NTD and Scm3 was calculated using the formulae:

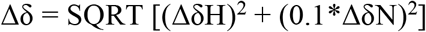

where, ΔδH is the difference in the amide proton cross-peak chemical shift and ΔδN is the difference in ^15^N chemical shift. The cut-off was decided according to the previously published method (34). In a similar fashion the CSP of ^15^N labeled Cse4 N-terminus with Scm3 (1:2) was determined. Corresponding blanks were recorded and all titration studies were performed thrice.

#### Relaxation studies

^15^N longitudinal (R1) and transverse relaxation rates (*R*2) of ^15^N Scm3 and ^15^N Scm3+N-terminus complex (1:2) were measured using CPMG delays: 25,50,100,200*,350,500,750*,1000ms 10,20*,40,60,80,100*,120,150 and 180 ms (* indicates repeat points) respectively. The intensities of the peaks were determined using Cara 1.8.4.2 analysis software and were fitted in a single exponential decay equation I(t) = A + Be−R_1/2_t in Sigmaplot 10.0 to obtain the R_1_ and R_2_ rates. The error (Δ*R*) in T_1_/T_2_ were calculated from the data-fitting errors as ΔT_1/2_/T_1/2_^2^ where Δ*T*_1/2_ is the standard deviation in the *T*_1/2_ fitting. The experiments were repeated twice. Error propagation for each of the residues was performed using the following equation:

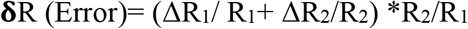

#### Labeling of Scm3 with (*S*-(1-oxyl-2,2,5,5-tetramethyl-2,5-dihydro-1H-pyrrol-3-yl)methyl methanesulfonothioate) and PRE-NMR studies

The labeling of Scm3 with the paramagnetic probe MTSL was done using the method described previously (35). Scm3(1mM) was diluted in labeling buffer (200 mM phosphate, pH7.4, 500mM NaCl) to a final concentration of 250-300 µM in 1 ml. The protein was desalted using the ZebaSpin™ (Thermofischer scientific) desalting column and incubated with 10X MTSL at room temperature for 15 min. Thereafter, 10X MTSL was further added to the protein mixture and kept on a rocker for 20hrs at 4°C, followed by incubation for 4hrs at room temperature. Excess and unligated MTSL was then removed by a step of buffer exchange using Amicon Ultra-15 centrifugal filter with the NMR buffer with no DTT (same as labeling buffer). The labeling of Scm3 with MTSL was verified by MALDI-TOF and Electron Spin Resonance. The ^1^H-^15^N HSQC spectra of paramagnetic and diamagnetic Scm3 were recorded at 283K and their intensity ratios were determined without and with Cse4 N-terminus using the CARA 1.8.4.2 software.

#### Diffusion Ordered Spectroscopy (DOSY)

The 1D and 2D experiments for Scm3, Cse4 N-terminus and their protein complex were carried out in 20 mM PB buffer at pH 7.4 at 25 °C at a protein concentration of 100 μM. First, 1D DOSY spectrum was obtained at 5% and 95% of the gradient strength (100% of gradient strength is equivalent to 48.4 G/cm) to standardize the gradient length and the diffusion delay. 2D DOSY were obtained using the diffusion delay of 150 ms and the gradient length of 3 ms for all samples. The DOSY spectra were obtained at 32 different gradient strengths varying from 2.42 to 45.98 G/cm with 64 scans at every increment. The DOSY data was then processed and analyzed by the Bruker Topspin 3.2 Software. The diffusion coefficients were obtained by fitting the intensity decay against the variable gradient strength. The hydrodynamic radii of the protein was calculated by the Stokes-Einstein equation,

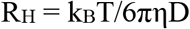

where R_H_ is the hydrodynamic radii, k_B_ is the Boltzmann constant, T is the absolute temperature, η is the viscosity of the PB buffer at 288K and D is the diffusion coefficient. The experiment was performed twice.

#### Pull down-Mass spectrometry

Bacterial lysate overexpressing yeast His-Scm3 was gel estimated for Scm3 by loading known concentrations of BSA and analyzing the comparative band intensities using the ImageJ software. 100µM of His-Scm3 (bait) was then bound to 200 µL Ni-NTA resin in two tubes followed by washes with TN buffers with increasing imidazole concentrations (25 and 50mM) to remove non-specifically bound impurities from the resin. Ni-NTA resin alone was used as control. In one tube, 500µL of 100µM of purified Cse4 N-terminus was added to the Scm3 bound resin for complexation and incubated for 1hr at RT while shaking. The flow-through was collected and the resin was washed with 2CV of TN buffer followed by addition of 10 mg/mL metaphase-arrested budding yeast lysate with a endogenous Scm3-HA tag. The yeast lysate was prepared as described previously (36, 37). The lysate was then incubated with both the resins (Scm3 bound and Scm3-N bound) overnight at 4°C while rotating. The next morning, the resins were washed with 2CV TN buffers with 25 and 50mM imidazole. The proteins (prey) bound to Ni-bound Scm3-His protein were eluted in 200 µL fractions with TN buffer containing 250 mM imidazole and stored at 4°C for Trypsin digestion and mass spectrometry.

#### Mass spectrometry

50µg of the samples were buffer exchanged and reduced with dithiothreitol (DTT) in denaturant and alkylated with sodium iodoacetamide (IAM) and digested with endopeptidase Trypsin. The resulting cleavage fragments were separated over a Phenomenex 100 Å Luna Omega 1.6 µm C18, 100 × 2.1 mm Column with a 155 min gradient of acetonitrile on a Shimadzu UPLC Nexera X2. The sample preparation was performed on two different days to check the reproducibility of data. Identification of the peaks were accomplished with an in-line Bruker MaXis II Q-TOF mass spectrometer and Biotools software version 3.2. Mass measurement accuracy of these experiments is expected to be within 50 ppm of the theoretical values. Each digested sample was queried against 88 different protein sequences from SGD (https://www.yeastgenome.org/) using Biotools 3.2 software. Carbamidomethyl of cysteine residues was selected as pre-modification and the number of mis-cleavage were set to maximum one. The percentage coverage at MS1 was obtained for all the proteins. All the samples were run in replicates of two.

## RESULTS

### Cse4 N-terminus is intrinsically disordered

NMR studies from our group revealed that the N-and C-terminus of Cse4 have two separate folding cores and behave as two distinct domains (20). Hence, we truncated the protein and selected the N-terminus of Cse4 for our studies. The protein was purified using affinity chromatography as described in Section 2 to obtain a pure band in 12% SDS-PAGE (Fig. 1A inset). *In silico* secondary structure prediction shows that the N-terminus is predominantly unstructured (20). The fact that the N-terminus could not be resolved in the Cryo-EM structure (2.7 Å) of the budding yeast specialized nucleosome further hints that it is disordered (38). The lack of structural information about the Cse4 tail is a bottleneck towards decoding the centromere composition and function.

**Fig. 1:**
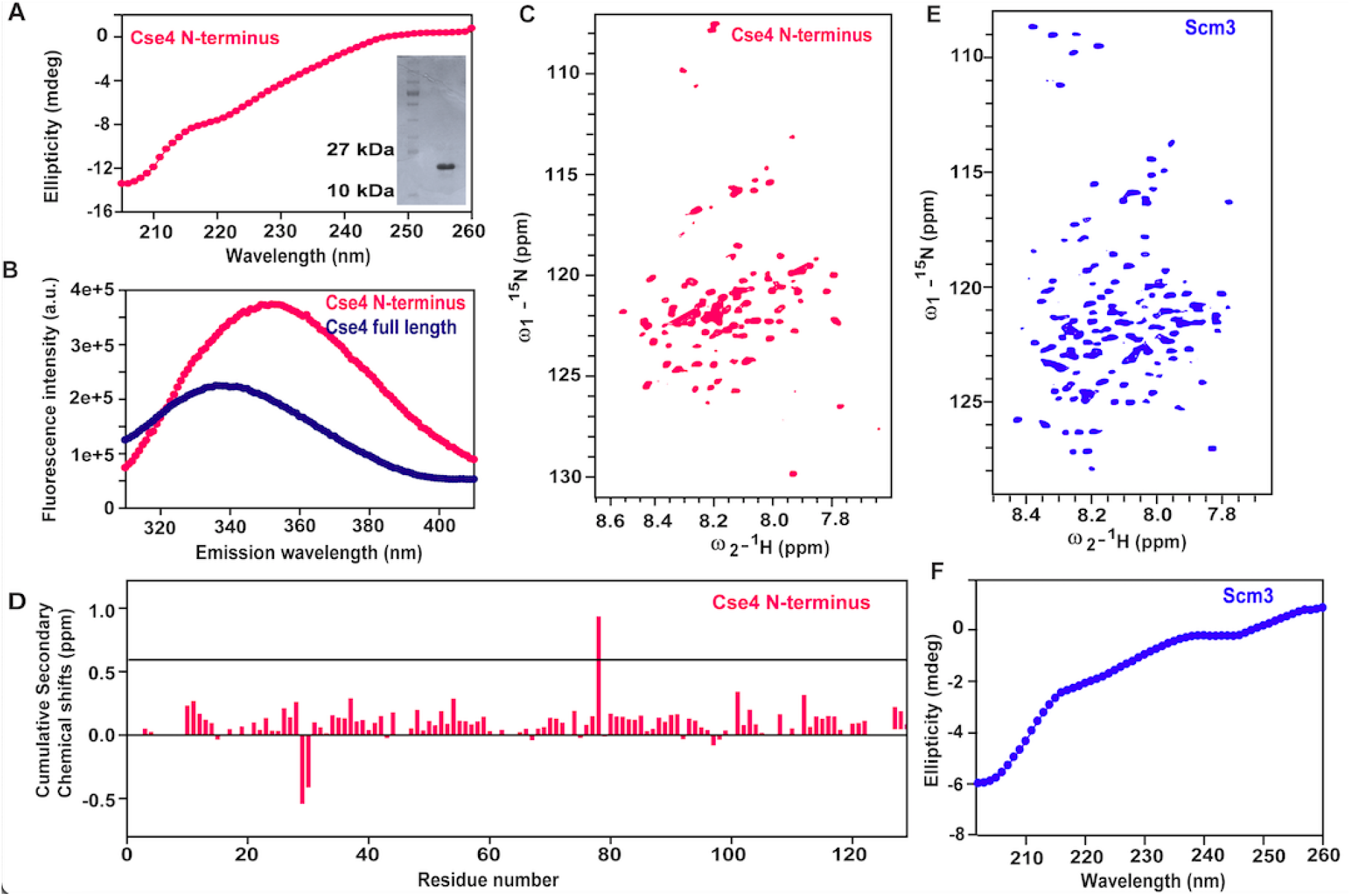
Cse4 N-terminus is intrinsically disordered: A) Far UV-CD spectra of Cse4 N-terminus. Inset: 12% SDS-PAGE of purified protein. B) Fluorescence spectra of Cse4 N-terminus and full length Cse4 (taken at different concentrations and times) C) ^1^H-^15^N HSQC spectra of ^15^N Cse4 N-terminus D) Cumulative Secondary Chemical Shifts of Cse4 N-terminus calculated from NMR backbone assignments, the black solid line indicates the cut-off above which a cluster of three or more residues are considered to be helical E) ^1^H-^15^N HSQC spectra of ^15^N Scm3 F) Far UV-CD spectra of Scm3.

First, we experimentally validated that the Cse4 N-tail is intrinsically disordered by the circular dichroism (CD) spectrum minimum at 200 nm and the fluorescence emission maximum at 350 nm (red shift) that indicate that the protein is in the random coil conformation (Fig. 1A, B). The low dispersion of the peaks along the proton (^1^H) dimension in the ^1^H-^15^N HSQC spectrum of the ^15^N labelled protein is also an indication of intrinsic disorder (Fig. 1C). Backbone resonance assignment of the protein was done using the standard 3D NMR experiments (HNCO, HN(CA)CO, HNCACB and HN(CO)CACB and HNN) at 298K and physiological pH (Supplementary Fig. 1, BioMagResBank (http://www.bmrb.wisc.edu) accession number 51203). This would be a foundational step to further decipher the structure, dynamics and interactions of the Cse4 N-terminus. Residue specific assignments could be successfully obtained for 103 out of 129 residues. Secondary structure propensities calculated from the backbone assignments showed that the N-terminus has no apparent secondary structure but has helix-formation tendencies (Fig. 1D) in some regions and which may lead to some closed topologies in the ensemble.

We have published the resonance assignment and the secondary structure propensities of Scm3 (BMRB accession number 27301) (27). The ^1^H-^15^N HSQC spectrum and the CD spectra minima indicate a random coil conformation in agreement with the previous observations (Fig.1E,F) (12, 27). Thus, both Cse4 N-terminus and Scm3 are classified as intrinsically disordered proteins. Bloom *et. al*. showed that the homology modelled Cse4 N-tail protrudes out of the nucleosome (39) (Supplementary Fig. 2A). Further, cryo-EM study of the specialized nucleosome revealed that the disordered tail makes minimum contacts with the DNA (38). This gives us a hint that the tail is available for post-translational modifications (PTMs) and interactions.

### Cse4 N-terminus interacts with Scm3 *in vitro* with a moderate affinity

The entire Cse4 N-terminus and partial regions of Scm3 have been deleted in the NMR structure of the reconstituted Cse4-Scm4-H4 complex (16). Given the ‘open conformation’ of the long N-tail of Cse4 in the Cse4-H4 heterodimer and the dynamic association of Scm3 at the centromere at all cell-cycle stages, it would be interesting to see whether the N-tail forms direct contacts with Scm3 and if it does, what could be the role of this interaction. For this, we probed an interaction between the full length proteins to provide a more comprehensive understanding of the interaction dynamics at the centromere. To test whether there is a direct interaction between Cse4 N-terminus and Scm3, purified recombinant proteins were run through a S200 column separately as well in a 1:1 equimolar ratio. A change in the elution profile of the 1:1 mixed proteins compared to those of the individual ones indicated complex formation. This was verified by running standards of appropriate molecular weights coinciding with those of Cse4 N-terminus, Scm3 and the complex. As expected, Cse4 N-terminus (MW∼15kDa) co-eluted with lysozyme (MW∼15kDa) while Scm3 (MW∼26kDa) co-eluted with carbonic anhydrase (MW∼30kDa) (not shown in figure). The elution profile of the complex (MW∼41kDa) was close to that of Ovalbumin (MW∼40kDa) confirming a direct interaction between Cse4 N-terminus and Scm3 *in vitro* (Fig. 2A). The elution peak of the complex when run on a 12% SDS-PAGE gel showed bands for both Cse4 N-terminus and Scm3 thereby confirming complexation between the two (Fig. 2B).

**Fig 2:**
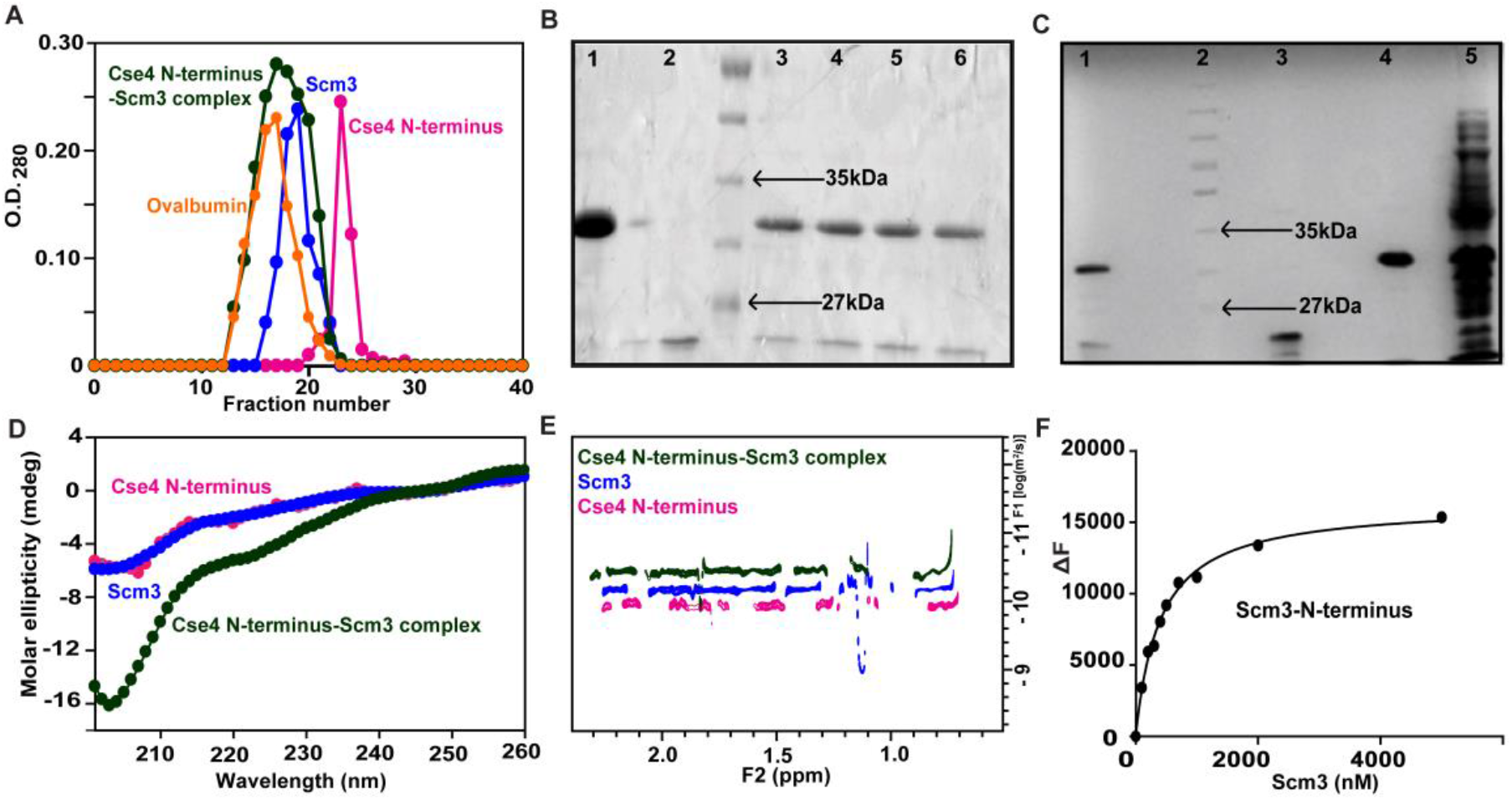
Cse4 N-terminus interacts with Scm3: A) Gel filtration elution profiles of Cse4 N-terminus (magenta), Scm3 (royal blue), Cse4 N-terminus-Scm3 complex (bottle green), Ovalbumin (orange) B) 12% SDS-PAGE gel indicating the bands corresponding to the peak co-eluted with Ovalbumin (3-6). Lane 1 and lane 2 denote pure Scm3 and Cse4 N-terminus protein respectively for reference and Lane 3 has the protein molecular weight marker C) Co-elution of N-terminus with Scm3 (Lane 1) in a pull-down assay. Purified Scm3 (Lane 4) and N-terminus (Lane 3) have been run for reference. Lane 5 indicates Scm3 lysate bound to resin used as bait. Lane 2 has the protein molecular weight marker D) Far-UV CD profiles of N-terminus (magenta), Scm3 (royal blue) and complex (bottle green) E) DOSY diffusion profile for Scm3 (royal blue), N-terminus (magenta) and Scm3-N-terminus complex (bottle green) F) Binding isotherm of N-terminus in presence of increasing concentrations of Scm3 for k_d_ determination.

Further, an *in vitro* pull-down assay was also performed to validate the interaction between Cse4 N-terminus and Scm3. Scm3-His bound to Ni-NTA resin was able to capture Cse4 N-terminus as both the protein bands were detected from the elute on a 12% SDS-PAGE gel (Fig. 2C) whereas no N-terminus band was seen with a resin-only control (data not shown). In contrast, no interaction was observed between Scm3 and H3 (Supplementary Fig. 2B) as Scm3 is a chaperone specific to Cse4 and does not interact with H3 (40). H3 was used as a negative control for the pull-down assay.

Additionally, far-UV CD studies also indicated an interaction between Cse4 N-terminus and Scm3 (Fig. 2D). Although no change was observed in the secondary structure content of the 1:1 complex in comparison to those of individual proteins, a change in ellipticity was seen. This indicates that while Cse4 N-terminus and Scm3 interact, they do not gain any significant secondary structure even after complexation. This observation presents an interesting mode of interaction between two intrinsically disordered proteins (IDPs) wherein there is no obvious gain in structure. When the same was repeated for a 1:1 Scm3-H3 complex, no change was observed (Supplementary Fig. 2C). The complexation between Cse4 N-terminus and Scm3 was also confirmed by Diffusion Ordered Spectroscopy (DOSY-NMR) based on the differential-diffusion rates of the 1:1 complex as compared to those of free proteins (Fig. 2E) and a significant increase in the Rh of the 1:1 complex (4.7±0.009 nm) is observed (Supplementary Fig. 2I). DOSY NMR along with the gel filtration assay suggests that Cse4 N-terminus and Scm3 form a stable complex *in vitro*.

The affinity of interaction between the two proteins was determined using fluorescence-based k_d_ assay as described in Section 2. When FITC-Cse4 N-terminus was titrated against varying concentrations of Scm3, a concentration-dependent increase in fluorescence of Cse4 N-terminus was observed (Supplementary Fig. 2F) which is also an indication of interaction. The change in the fluorescence (ΔF) was fitted in a binding isotherm and a k_d_ value of 400±10.2 nM was obtained (Fig. 2F). The results therefore indicate that Cse4 N-terminus interacts with Scm3 with a moderate affinity. We were unable to obtain the C-terminus of Cse4 *in vitro* in a monomeric state and hence full-length Cse4 was used as a positive control for the assay. The k_d_ of Cse4-Scm3 interaction was calculated to be 579±5 nM (Supplementary Fig. 2F). H3 was used as a negative control and as expected, in presence of increasing concentrations of H3, a negligible change in the fluorescence of FITC-Scm3 was observed, indicating an absence of interaction (Supplementary Fig. 2G). Previously, Dechassa *et. al*. demonstrated that Scm3 interacts with the (Cse4/H4)2 tetramer with a nanomolar affinity (∼18nM) (41). Notably, the k_d_ values support that Scm3 is a chaperone of Cse4-H4 rather than of Cse4 alone. Furthermore, the affinity did not change in the presence of the N-terminus (41), indicating that it plays no direct role in centromere deposition. This is supported by the observation that Cse4 exists in an ‘open conformation’ only in the presence of H4, allowing the N-terminus to freely interact with other proteins (21). Additionally, α-synuclein, a known IDP was also used as a negative control to validate the specificity of the interaction between Cse4 N-terminus and Scm3 (Supplementary Fig. 2H). Therefore, our results suggest that Cse4 N-terminus and Scm3 specifically bind to each other and form a stable complex *in vitro*.

### The interaction between Cse4 N-terminus and Scm3 is driven by electrostatics

We investigated what forces drive the tail-chaperone protein complex in order to gain mechanistic insights into their interaction. Both Cse4 N-terminus and Scm3 are intrinsically disordered proteins with multiple acidic and basic patches (Fig. 3A,B) (42). Hence, one could assume that these regions may contribute towards an interaction between the two. To test whether the interaction is driven by electrostatics, the affinity of interaction (kd) between the two proteins was determined at two different salt concentrations: 50 and 500 mM KCl. NaCl could not be used for the assays due to the instability of the proteins at low NaCl concentrations. Interestingly, the k_d_ value decreased in the presence of 500 mM KCl when compared to 50 mM KCl (1.5 times lesser) (Fig. 3C) indicating screening of charges on the protein by the salt ions thereby rendering them unable for interactions. The results, therefore, indicate that the interaction between Cse4 N-terminus and Scm3 is governed by electrostatics with opposite charges on both the proteins interacting with each other (Fig. 3F). This was further verified by probing the contribution of hydrophobic forces in the interaction using an ANS-based assay. The change in the fluorescence spectra of ANS-labelled Cse4 N-terminus with increasing Scm3 concentrations was random, indicating that hydrophobic interactions had no role in Cse4 N-terminus-Scm3 binding (Fig. 3D). Electrostatic interactions between oppositely charged histone tails and chaperone has been reported for H1 and ProTα. The large and opposite net charges of ProTα (−44) and H1 (+53) have a significant contribution to binding between the two IDPs (43).

**Fig 3:**
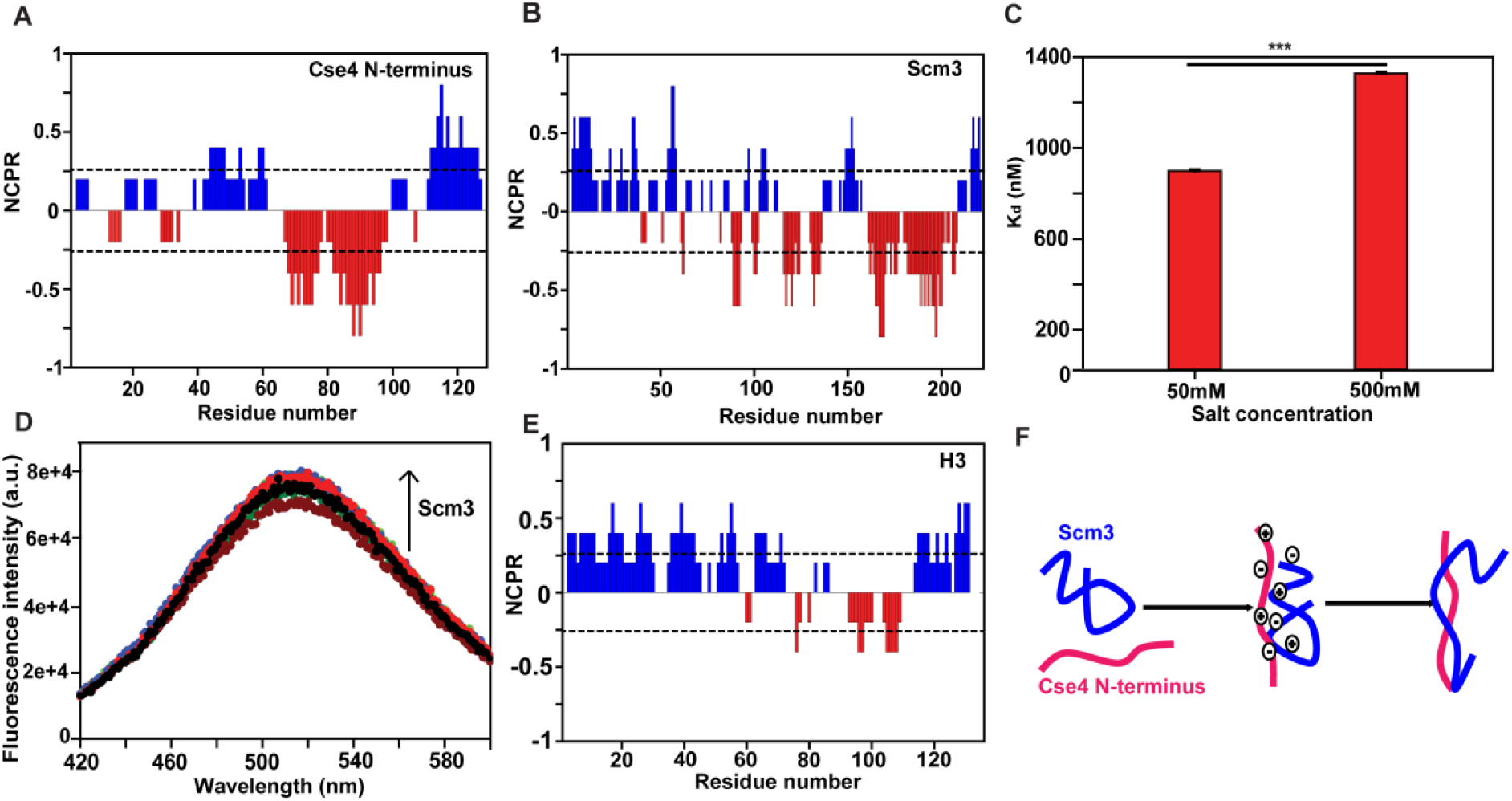
The interaction between Scm3 and N-terminus is driven by electrostatics: A) Charge distribution of N-terminus as determined using CIDER. Red bars denotes negative charges while blue bars denote positive charges B) Charge distribution of Scm3 as determined using CIDER. Red bar denotes negative charges while blue bar denotes positive charges C) Comparison of k_d_ between 100 nM FITC-N-terminus and Scm3 (100-5000 nM) in presence of 50 and 500 mM KCl D) ANS binding profile of N-terminus in absence or presence of increasing concentrations of Scm3. The upward arrow indicates increasing concentrations of Scm3 E) Charge distribution of H3 as determined using CIDER F) Proposed mode of binding between N-terminus (magenta) and Scm3 (royal blue).

To check the specificity of the IDP-IDP interaction, we explored if Scm3 interacts with α-synuclein, another IDP, close in size to Cse4 N-terminus. However, these two proteins do not appear to interact because changes in the fluorescence intensity of FITC-Scm3 upon addition of α-synuclein in increasing concentrations are non-specific (Supplementary Fig. 2H). Scm3 also fails to interact with H3 as previously mentioned. Furthermore, mixing H3 and Scm3 in a 1:1 ratio resulted in a white cloudy precipitate (Supplementary Fig. 2A) with no visible droplet formation, making us believe that H3 and Scm3 are incompatible with one another. The uniqueness of the specific IDP-IDP interaction between N-terminus and Scm3 can be attributed to the charge patterning (distribution of positively and negatively charged residues) with the opposite charges on both proteins interacting with each other. While the acidic and basic patches in N-terminus are more distinct, the charge patterning in H3 (Fig. 3E) and α-synuclein are less pronounced. The same is reflected quantitatively in the ‘Ƙ’ (kappa) parameter obtained from the CIDER analysis. As Ƙ increases from 0 to 1, the sequences become less well mixed in terms of positive and negative residues. The Ƙ values of the protein of interest are summarized in Table 2. Interestingly, both N-terminus and Scm3 have Ƙ values that are closer to 0 than to 1, indicating that the IDPs have a well-mixed charge composition, i.e. the intra-chain electrostatic forces are counter-balanced, and thus have no bias towards a particular conformation. Instead, they may sample a number of unbiased random coil conformations, as expected for IDPs (44). To assess if these Ƙ values are substantially more ‘blocky’ (charge-segregated) than expected by random chance, we computed the Z-score associated with the sequence Ƙ value compared to a null distribution of randomly shuffled sequences with the same composition (45). While Cse4 N-terminus and Scm3 were almost three standard deviations from the mean (Z-scores of 2.97 and 2.72, respectively) H3 and α-synuclein were both very close to the expected value (Z scores of 0.28 and –0.01, respectively). Taken together, our analysis supports a model in which Cse4 N-terminus and Scm3 contain charge-segregated subregions that drive interaction.

**Table 2:**
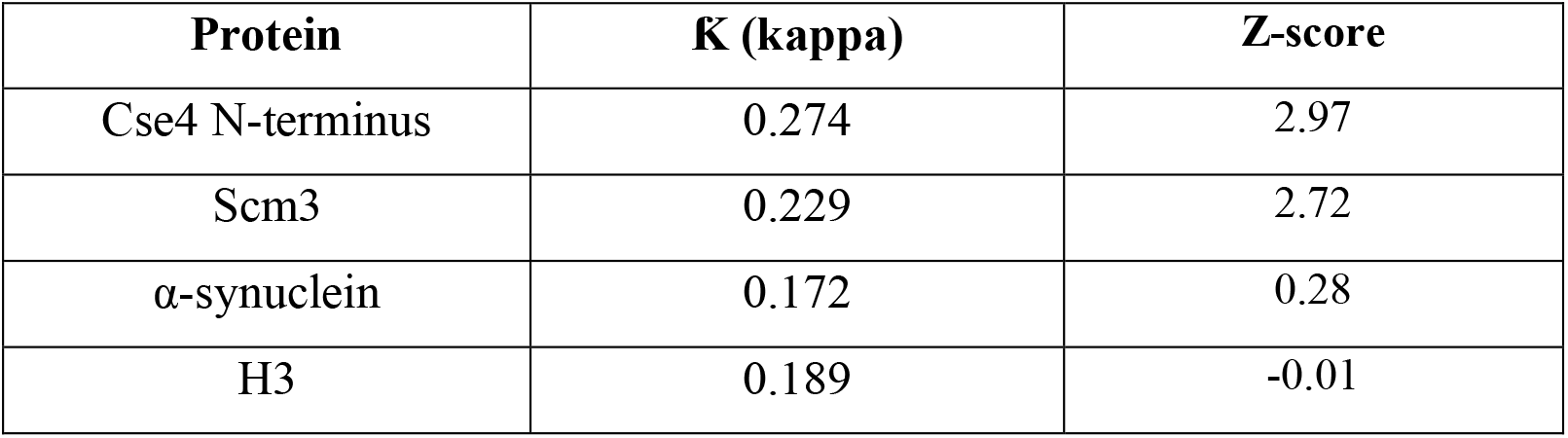
Ƙ values of relevant proteins.

| **Protein** | **K (kappa)** | **Z-score** |
| --- | --- | --- |
| Cse4 N-terminus | 0.274 | 2.97 |
| Scm3 | 0.229 | 2.72 |
| α-synuclein | 0.172 | 0.28 |
| H3 | 0.189 | -0.01 |

**Table 3:** Intrinsic disorder content in KT proteins as predicted from IUPred2A.

| KT protein | IDR/Total | % |
| --- | --- | --- |
| Cbf1 | 260/340 | 76.47059 |
| Cnn1 | 250/360 | 69.44444 |
| Rad52 | 320/480 | 66.66667 |
| Scm3 | 140/223 | 62.78027 |
| Mps1 | 420/680 | 61.76471 |
| Psh1 | 220/400 | 55 |
| Mif2 | 270/520 | 51.92308 |
| Cse4 | 110/229 | 48.03493 |
| Skp1 | 70/190 | 36.84211 |
| Okp1 | 110/410 | 26.82927 |
| Dsn1 | 158/576 | 27.20000 |
| Stu2 | 220/840 | 26.19048 |
| Mre11 | 180/700 | 25.71429 |
| Ndc80 | 120/680 | 17.64706 |
| Ctf19 | 60/365 | 16.43836 |
| Ame1 | 50/318 | 15.72327 |
| Mcm21 | 40/360 | 11.11111 |
| Chl4 | 10/450 | 2.222222 |

### Cse4 N-terminus and Scm3 interact with each other at multiple sites and the latter is stabilized by the former

To map the residues that are involved in interaction on both Cse4 N-terminus and Scm3, we employed NMR spectroscopy. The ^15^N-^1^H heteronuclear single quantum coherence (HSQC) spectrum of Scm3 has a low ^1^H dispersion consistent with its disordered nature (Supplementary Fig.3A) (27). The resonance assignment of full length Scm3 has been previously published by our group and was used for residue mapping (27). Firstly,^15^N-Scm3 was titrated without and with equimolar ratio of unlabeled Cse4 N-terminus to check the chemical shift perturbations caused on Scm3 by Cse4 N-terminus. Interestingly, it was observed that the ^1^H chemical shift dispersion of 15N Scm3 remained unchanged even in the presence of Cse4 N-terminus, demonstrating that the protein complex does not gain any significant structure. This is in line with our CD findings (Fig. 2D). However, quantifiable changes in the chemical shifts of ^15^N Scm3 were observed in the presence of N-terminus (Supplementary Fig. 3A) and were used for residue-wise mapping. The residues with non-distinct peak centres were excluded from the analysis. Significant chemical shift perturbations (CSPs) were observed throughout the sequence (Figure 4A). Consistent results were obtained when the experiment was repeated thrice. Representative chemical shift perturbations for few residues have been shown as inset (Fig. 4A). Some residues such as K139 and E53 (inset) whose peak centres are merged in Scm3 alone, appear to be well-resolved in the complex indicating that N-terminus causes local structural changes in Scm3 on interaction.

**Fig 4:**
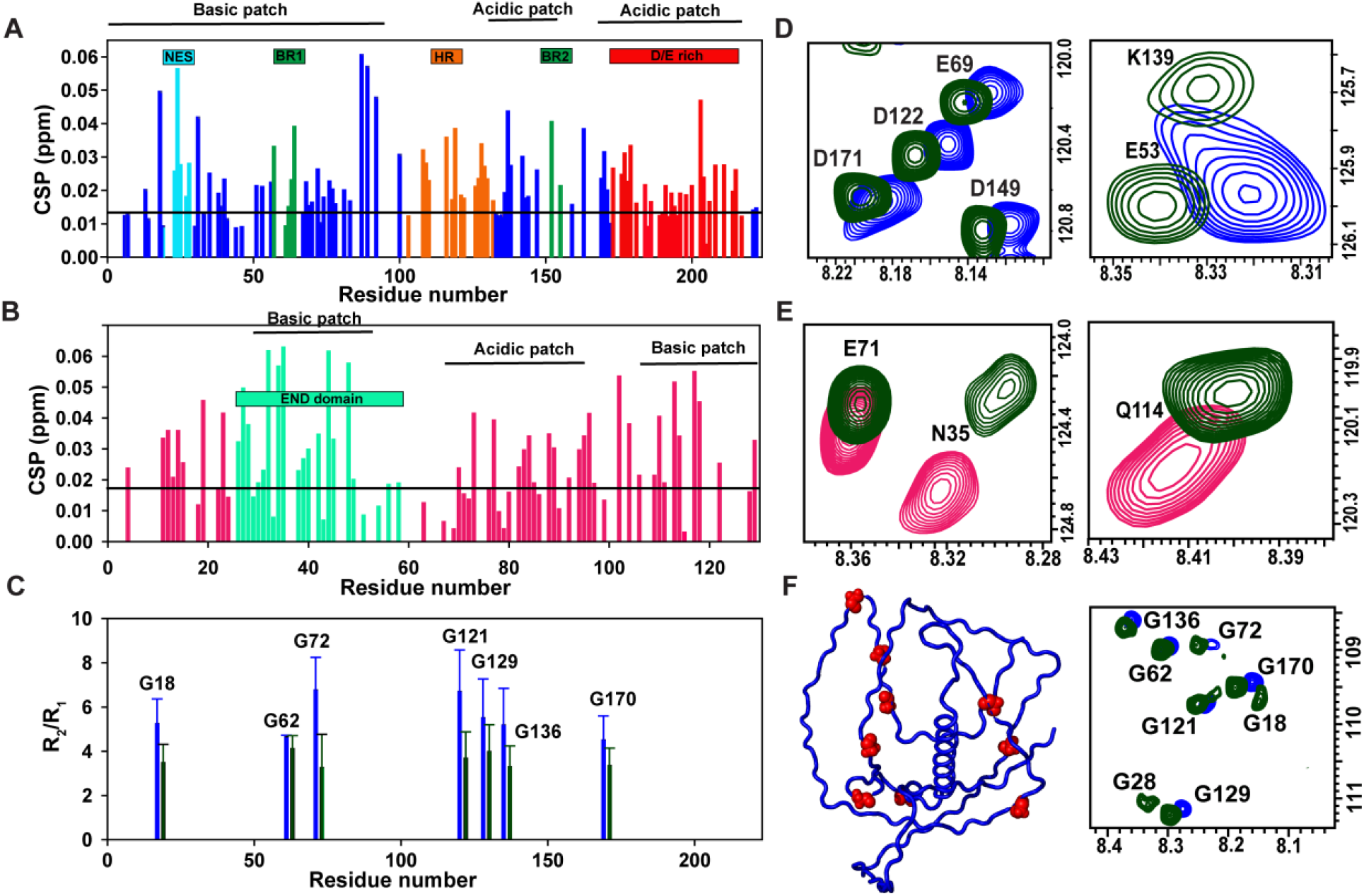
The interaction between Scm3 and N-terminus is multisite: A) Left: Residue-wise CSP changes of ^15^N Scm3 in presence of two-fold molar excess of N-terminus. Right: Representative CSP of residues in ^15^N Scm3 HSQC spectra in the presence of N-terminus B) Left: Residue-wise CSP changes of ^15^N N-terminus in presence of two-fold molar excess of Scm3. Right: Representative CSP of residues in ^15^N N-terminus HSQC spectra in the presence of Scm3. Annotated domains in both the proteins have been highlighted in different colours. C) Left: R_2_/ R_1_ of Glycine residues of ^15^N Scm3 alone (royal blue) and in presence of N-terminus (bottle green). Right: Glycine region in the HSQC spectra of ^15^N Scm3 (royal blue) and in complex with N-terminus (bottle green).

The result therefore indicates that Scm3 has multiple binding sites for Cse4 N-terminus. As mentioned earlier, the N-terminal Cse4 binding domain (CBD) of Scm3 undergoes a disorder-to-order transition upon binding to Cse4 HFD (12) while our findings suggest that Scm3 does not fold upon binding to N-terminus. This highlights the promiscuous behaviour an IDP can adopt in the presence of different binding partners (46). Additionally, the N-terminus of Scm3 would be available for interaction with the Cse4 N-terminus only after its deposition at the centromere. This would imply that Scm3 interacts with N-terminus after the deposition of Cse4 at the centromere.

Scm3 binds to centromeric (CEN) DNA through a predicted DNA binding region (DBR) (1-103) (47). We found that Scm3 has a very high affinity for CEN3 DNA (3.4±1 nM) (Supplementary fig. 3E,F) when compared to N-terminus (400±10 nM). Hence, to check whether CEN-bound Scm3 can bind to N-terminus, we performed a competition assay wherein a saturated complex of FITC-Scm3 and CEN3-DNA was made and N-terminus was added in increasing concentration. No significant change in the fluorescence intensity was seen which indicates that N-terminus cannot associate with CEN3-bound Scm3 (Supplementary fig. 3G). Besides, the DBR in Scm3 (∼1-103) is also involved in an interaction with Cse4 N-terminus (Fig. 4A) and it would have to be free from CEN3-DNA for doing the same. In fact, Scm3 is in a dynamic exchange at the centromere which will allow it to associate with several proteins in a cell cycle stage-specific manner (19). Further, Cryo-EM and biophysical studies of the budding yeast CENH3 nucleosomes revealed that the N-terminus makes very few contacts with the CEN DNA (38, 48). This means that the N-terminus is also free from the CEN DNA to interact with enzymes, remodelers and other proteins-one of them being Scm3. In short, Scm3 and N-terminus interaction is free from the influence of the CEN DNA.

The binding regions on ^15^N Cse4 N-terminus involved in interaction with Scm3 were also identified by NMR (Supplementary Fig 3B). Remarkably, even in the presence of equimolar concentrations of unlabeled Scm3, no change in the chemical shift dispersion along the ^1^H dimension was detected, demonstrating the absence of gain of any form of structure. However, noticeable changes in chemical shifts of N-terminus were detected with addition of Scm3. The residue mapping of the ^15^N Cse4 N-terminus peaks was done from the assigned amino acids (Supplementary Fig. 1). Observed CSPs were plotted as a function of the residue number to map interacting sites on N-terminus (Fig. 4B). Consistent results were obtained when the experiment was repeated thrice. Representative chemical shift perturbations for few residues have been shown as inset in Fig. 4B right panel. An overall change in CSPs was obtained for N-terminus except for the residues ∼50-70. Markedly, residues 50-60 form a part of the Essential domain (END domain) of N-terminus that is directly involved in an interaction with the Okp1 core domain (22, 23). Overall, NMR studies of both N-terminus and Scm3 indicate a multisite mode of interaction with no considerable gain in secondary structure. This multisite interaction is consistent with the inference that the interaction is driven by electrostatics since the proteins are seen to have discrete acidic and basic patches along the sequence. This type of IDP-IDP interaction is reminiscent of the one adopted by the proteins 4.1G and NuMA (49).

Further, to check the effect of the Cse4 N-terminus on Scm3 dynamics, transverse relaxation rates (R2) of non-overlapping glycine residues of ^15^N Scm3 were measured in absence and presence of the Cse4 N-terminus. Due to large overlapping regions in the Scm3 spectra, accurate intensity measurements were difficult for all the residues. Scm3 has nine well-resolved glycine residues that span across its sequence and hence measuring the R2 rates of the glycine residues should give a quantitative picture of the interaction dynamics of Scm3 with the Cse4 N-terminus. Interestingly, there was a significant decrease in the R2 rates (Average values: Scm3-12.7± 1.8 s^-1^ and Scm3N complex-9.81±1.5 s^-1^) and increase in R1 (Average values: Scm3-2.3±0.35 s^-1^ and Scm3N complex-2.7±0.38 s^-1^) rates of all the glycine residues in the complex (Supplementary Fig. 3&D). Additionally, it was observed that the complex has a better signal-to-noise (S/N) ratio in comparison to that of ^15^N Scm3 alone (for instance, G28 and G18 have reappeared in the complex; See Fig. 4C, right panel) and hence the decrease in the R2 rates could be due to a decrease in the exchange component (R_ex_) of Scm3 in the presence of the N-terminus. Indeed, upon calculation of R2/ R1 (∼R_ex_) ratios of the glycine residues in Scm3 in presence of the N-terminus, a significant decrease was observed when compared to that of free Scm3 (Average values: Scm3-5.6±1.6 and Scm3N complex-3.6±1) (Fig. 4C). Thus, N-terminus suppresses exchange in Scm3 and this would implicate its stabilization at the centromere and we speculate that this stabilization may be important in the context of KT assembly. The NMR ensemble of Scm3 predicted by CS23D2.0 (Supplementary Fig. 2E) can be seen to have multiple conformations and the N-terminus possibly reduces the conformational heterogeneity of Scm3 to an extent. Notably, Scm3 does not gain any substantial secondary structure and hence may still retain its potential to bind to multiple proteins at the KT. Stabilization without folding is a mechanism of IDP interaction with its binding partner which has not been explored much (26, 50).

### N-terminus disrupts intramolecular contacts of Scm3 to stabilize it

Scm3 has one Cysteine (C41) residue which was utilized for paramagnetic labelling via S-(1-oxyl-2,2,5,5-tetramethyl-2,5-dihydro-1H-pyrrol-3-yl)methyl methanesulfonothioate (MTSL) in order to probe long-range interactions within Scm3. The labelling was confirmed by Electron Paramagnetic Resonance (EPR) and Matrix Assisted Laser Desorption / Ionization (MALDI) experiments (data not shown). The absence of C41 peak (closest to the paramagnetic tag) in the -15N-^1^H HSQC spectra of Scm3-MTSL confirmed the labelling (Supplementary Fig. 4A). Apart from the C41 where the labelling was done, and its immediate vicinity, calculation of the ratio of the peak intensities in the paramagnetic Scm3 with respect to the diamagnetic Scm3 (Iox/Ired) revealed several regions with decreased intensity (Supplementary Fig.4C). The calculations were done after acquiring a high-resolution ^15^N-^1^H HSQC spectrum of Scm3 by increasing the td points and acquisition times. Regions of high overlap were omitted from the analysis to avoid erroneous conclusions. MTSL was able to exert its paramagnetic influence on various regions of the protein other than its labelling position which shows that it has extensive transient long-range intramolecular contacts. It was observed that a number of peaks apart from C41 were missing in Scm3-MTSL notably residues 18,19,27, 28, 51, 53, 72, 76, 82, 92, 106, 118, 127, 149 and 167 which means these residues come in close contact with C41 (<12 Å). Majority of the Iox/Ired ratios are in the range of 0.5-0.6, indicating that most of the residues in the protein are in proximity of C41 (∼25 Å) (51) and that Scm3 is not completely extended. This indicates that Scm3 forms widespread long-range intramolecular interactions in its native state, and despite being intrinsically disordered, has some compact topologies in the ensemble of conformations. The PRE effect observed in Scm3 is similar to that observed for the intrinsically disordered C-terminus of the H2A/H2B chaperone Nucleoplasmin (Npm) indicating the presence of extensive intramolecular interactions. The long-range electrostatic interactions between the acidic and basic tracts of Npm regulates its interaction with histones (52).

To further corroborate our claim, we recorded the ^15^N-^1^H HSQC spectra of the denatured Scm3 in 8M urea (Supplementary Fig.4B). Clearly, the peaks in denatured Scm3 were significantly shifted in comparison to native Scm3 which further indicates that Scm3 has transient intramolecular interactions in its native state that are disrupted by 8M urea. Moreover, the NMR ensemble predicted for Scm3 also shows that Scm3 is a compact IDP with several intramolecular contacts (Supplementary Fig.2E). Interestingly, the residues 90-115 sample an alpha helical conformation and are part of the conserved heptad repeat in Scm3 (53). This serves as a good validation of the prediction algorithm. Further, it is the same region that assumes a helical structure in the presence of Cse4 HFD (54). The theoretical Rh calculated for Scm3 is 4.8 nm while the experimental Rh as determined by DOSY-NMR was found to be 3.8±0.025 nm which is also an indication that Scm3 is a compact IDP (Supplementary Fig.4E).

Interestingly, on addition of the N-terminus in ^15^N Scm3-MTSL, the peak intensities increase overall and some peaks (such as S39 and E118) that had disappeared under the paramagnetic influence of MTSL, reappeared in presence of the N-terminus (Supplementary Fig. 4D and G). This shows that the N-terminus breaks the extensive intramolecular contacts of Scm3 to some extent. Thus, from the NMR data, it can be inferred that the N-terminus interacts with Scm3 and suppresses exchange by disrupting intramolecular contacts in Scm3, thereby making a stable complex (Supplementary Fig. 4F). We suspect that it could also be an auto-regulatory mechanism wherein Scm3 is in an accessible conformation only upon binding to the N-terminus. Recently, an auto-inhibitory mechanism has been proposed for another essential inner KT protein, Mif2 whose disordered N-terminal tail exists in an inaccessible conformation and only opens up when it binds to the Cse4 nucleosome (55).

### Scm3 strengthens the interaction between Cse4 N-terminus and Ame1/Okp1

The N-terminal tail of Cse4, specifically the END domain, has been known to interact with the AO heterodimer (22, 23, 56). A truncated peptide of the N-terminus (33-110) binds to AO with a high affinity of 129±39 nM (22). Further, the core domain of Okp1 is involved in the direct interaction with the END domain of the N-terminus (23). We found that the dissociation constant (kd) of the interaction between the whole N-terminus (1-129) with AO is 44.7 ± 4 nM (Supplementary Fig. 6A). This is in agreement with the previous report which states that the affinity of this interaction increases with the increase in the peptide length of the N-terminus (22). The result indicates that there might be regions in the N-terminal tail, other than the END domain which may be involved in strengthening the interaction with AO, and the long tail may be a key player in mediating/strengthening other interactions as well. It has been shown that R37Me and K49Ac PTM marks on the Cse4 N-tail reduces its affinity for AO (22). To check if these modifications can alter the charge patterning on the N-terminus, we substituted both R37 and K49 to L (Leucine) as leucine mimics the hydrophobicity provided by the methylation and acetylation marks. There is an obvious decrease in charges at the END domain after leucine substitution (Supplementary Fig. 5) and this might cause a disruption in the interaction of the Cse4 N-terminus and Scm3 as their binding is largely driven by electrostatics (Fig. 3C). This leaves us with the question: Does Cse4 tail-chaperone interaction influence tail-AO binding?

Interestingly, a pull-down mass spectrometry (MS) assay with Scm3 as the bait detected both the Ame1 and Okp1 proteins (Fig.5G). Apart from AO, Scm3 also pulls down Cep3 (57), components of Ndc80 complex, components of the DASH complex, outer KT protein Chl4, mitotic checkpoint protein Bub3, DNA repair protein Rad50 and histone acetyltransferase Rtt109 according to their coverage percentages. Some of these coverages have been represented as heat maps in Fig. 5G. Moreover, significant coverages were not seen for H3 and Mif2 but were observed for Ndc10, Cse4 and H4 when compared to control thus validating our MS results (18). Therefore, the role of Scm3 as an important and versatile centromeric protein apart from being a specific histone chaperone is reinforced in our study.

**Fig 5:**
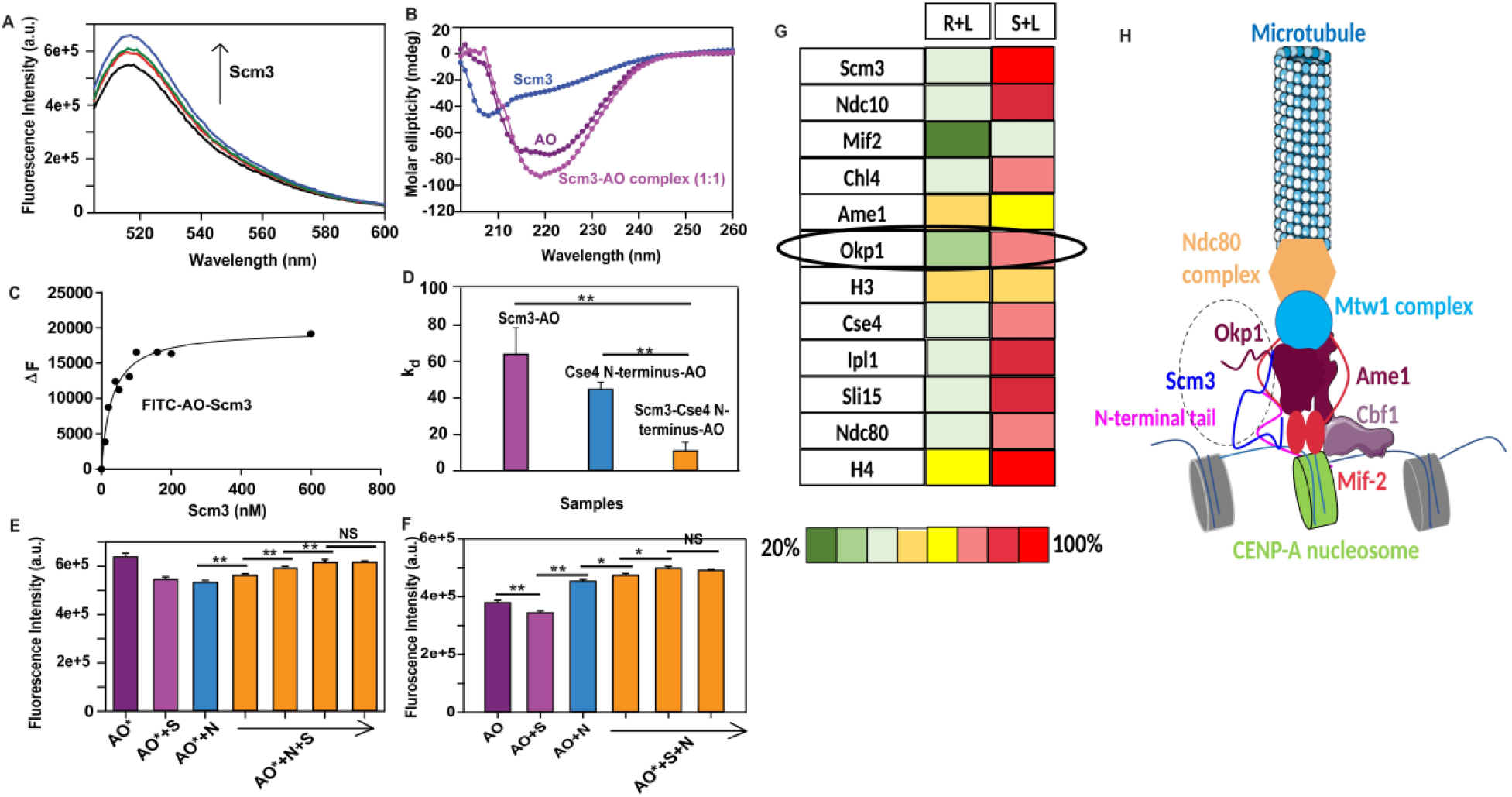
Scm3 interacts with AO: A) Fluorescence spectra of 50nM FITC-AO in absence (black) and presence of increasing concentrations of Scm3 (red, green and blue). B) Far-UV CD profiles of AO (purple), Scm3 (royal blue) and AO-Scm3 complex (dark pink) C) Binding isotherm of 50nM FITC-AO in absence and presence of increasing concentrations of Scm3 (5-500 nM) for k_d_ determination D) Bar graph of comparison of k_d_ of Scm3-AO, N-terminus-AO and Scm3-N-terminus-AO E) Assay of N-terminus-Scm3-AO: Purple bar represents the fluorescence intensity maxima of 50 nM FITC-AO denoted as AO (purple), 50 nM FITC-AO+ 50 nM Scm3 complex denoted as AO+S (dark pink), 50 nM FITC-AO+50 nM N-terminus complex denoted as AO+N (blue), 50 nM FITC-AO+ 50 nM N-terminus complex in presence of increasing concentrations of Scm3 denoted as AO+N+S (orange) F) Assay of N-terminus-Scm3-AO: Purple bar represents the fluorescence intensity maxima of 50 nM FITC-AO denoted as AO (purple), 50 nM FITC-AO+ 50 nM Scm3 complex denoted as AO+S (dark pink), 50 nM FITC-AO+50 nM N-terminus complex denoted as AO+N (blue), 50 nM FITC-AO+ 50 nM Scm3 complex in presence of increasing concentrations of N-terminus denoted as AO+S+N (orange). For E and F, error bars were added based on three readings acquired on the same samples. The experiment was repeated thrice and consistent results were obtained G) Heat map of the proteins interacting with Scm3 created based on the coverage percentages in control and test sample. R+L denotes resin+lysate and S+L denotes Scm3+lysate H) Schematic of the kinetochore depicting the synergy between AO-Scm3-N-terminus marked inside the dotted circle.

Now, since 1) the N-terminus interacts with AO and also forms a stable complex with Scm3 with no apparent gain in structure 2) AO is pulled down by Scm3 as detected in mass spectrometry 3) PTMs that control the N-terminus-AO interaction may also affect the N-terminus and Scm3 interaction 4) Scm3 recruits and maintains essential inner KT proteins such as Mif-2 and Ndc10 (18), we investigated whether there is a direct interaction between Scm3 and AO. It was observed that addition of Scm3 caused a change in the fluorescence intensity of FITC-AO in a concentration dependent manner (Fig. 5A) suggesting that there is an interaction between the two. This is also reflected in the far-UV CD experiments wherein a clear change in ellipticity and slight reduction in helical content of AO is observed in the presence of Scm3 (Fig. 5B). The k_d_ between Scm3 and AO was calculated to be 64 ± 14.5 nM which indicates a strong association between the two (Fig. 5C).

Since Scm3 interacts with both the N-terminus and AO, we checked how the presence of Scm3 can affect the binding between the two. For this, we titrated FITC-AO with increasing concentrations of 1:1 complex of N-terminus and Scm3, and found that the binding affinity increased significantly (∼3 folds) (Fig. 5D, Supplementary Fig. 6B). There appears to be a synergistic effect of Scm3 on N-terminus-AO binding. Further, to verify that the three proteins are involved in a synergistic association, a 1:1 AO-Scm3 complex was made in which both AO and Scm3 were labelled with FITC in two different experiments followed by a titration with increasing concentrations of N-terminus. A concentration-dependent increase in fluorescence intensity was observed followed by saturation demonstrating that the N-terminus can bind to both the proteins in the complex (Fig. 5F). The same was observed when a 1:1 complex of FITC-AO and N-terminus was made and titrated against Scm3 (Fig. 5E). Similarly, when 1:1 complex of Scm3 and N-terminus in which both Scm3 and N-terminus were labelled with FITC in two different experiments, a concentration-dependent increment in the fluorescence intensity was seen when AO was added signifying that both the proteins can interact with AO (Supplementary Fig. 6C, D). A schematic of KT representing the synergistic association has been shown in Fig. 5H. Hence, the lack of a defined structure in the Cse4 N-tail and chaperone Scm3 complex promotes association with a third protein at the centromere; forming a high-affinity complex.

## Discussion

The available structural studies for yeast CENH3 have only focused on the C-terminus HFD of Cse4 and the CBD of Scm3 leaving gaps in the interaction dynamics of Cse4 and Scm3. We show that the N-tail of Cse4 interacts with its chaperone Scm3 using multiple *in vitro* experiments such as Gel filtration assay, Pull-down assay, NMR and fluorescence. While previously it was reported that the two proteins do not interact with each other in a yeast two-hybrid assay (Y2H) (53), we observed a relatively strong interaction between the two (k_d_ = 400 nM). Interestingly, we were unable to detect protein expression for Cse4 N-terminus in budding yeast (*in vivo*), while the C-terminus could be successfully expressed under similar conditions (data not shown). This could be due to the rapid intracellular degradation of the N-tail, which is why the interaction was not captured in Y2H. Further, it has been speculated that the N-terminal peptide on itself is not stable and perhaps must be physically linked with the HFD for function *in vivo* (8).

There are two possibilities of when the tail-chaperone interaction may happen: Firstly, it has been shown that Scm3 binds to Cse4/H4 tetramer with a much stronger k_d_ of 18nM and the presence of the N-terminus does not affect the affinity (58). This suggests that Scm3 may associate with the N-terminus only after Cse4 is deposited at the centromere. Scm3 dislodges from Cse4/H4 to allow nucleosome assembly, following which it experiences a helical-to-disorder transition to return to its native state to interact with the N-terminus. Scm3 is in a continuous dynamic exchange at the centromere where old molecules are replaced with new ones every ∼5 minutes (19). This would allow Scm3 to associate with multiple proteins in a cell cycle stage-specific manner. Hence, the second possibility is that a different molecule of Scm3 (which is not DNA bound, Fig. 3.6G) interacts with the Cse4 N-terminus at the centromere.

**Fig. 6:**
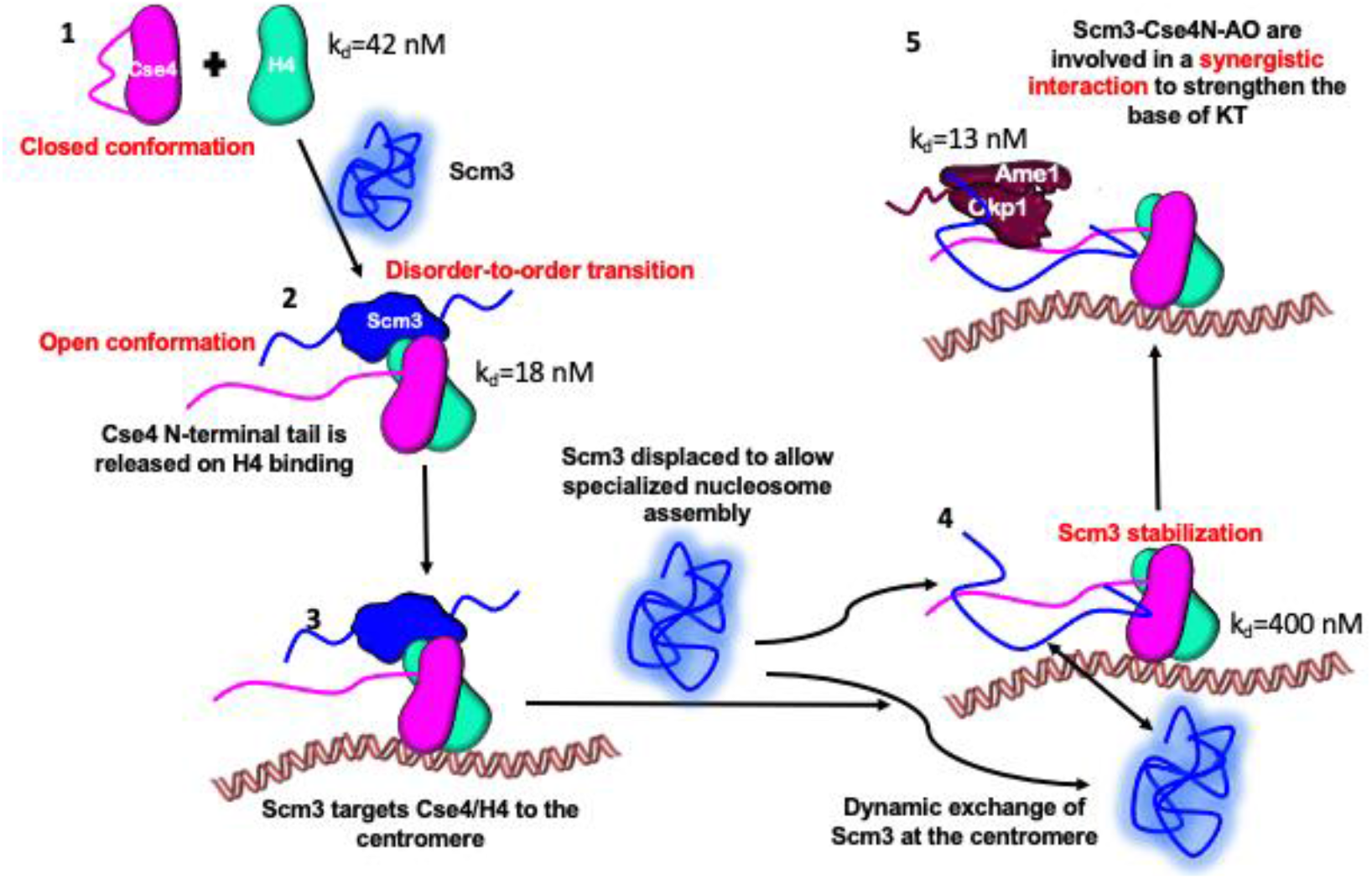
Proposed model: 1) Cse4 is in a closed conformation in native form 2) Upon binding to H4, N-terminus is free and Scm3 targets the dimer to centromere (3) after which it is dislodged to allow nucleosome assembly. Scm3 regains its native disordered form with extensive intramolecular contacts. It binds to the Cse4 N-terminus and the latter breaks intramolecular contacts and suppresses exchange in Scm3 to stabilize it (4). Scm3 is now in a more accessible conformation and makes contacts with AO during KT assembly. Scm3 strengthens the interaction of AO with N-terminus by forming a synergistic complex to stabilize the KT (5).

We have established that both the proteins are intrinsically disordered and form a stable protein complex *in vitro* with no significant gain in structure. Further, the interaction is multisite between the opposite charges on the proteins. This provides a moderate binding affinity to the protein complex as opposed to the weak and transient binding expected for IDPs. The complex is driven by electrostatic forces and the specificity is encoded in the charge patterning. Electrostatic forces have a significant long-range contribution in proteins or protein complexes as charge–charge interactions can be strong even at long distances (e.g., 5–10 Å) (59). This is noteworthy as no definite structure-formation accompanying binding can be easily supported by long-range interactions. This type of an interaction mechanism may provide a channel for crosstalk between multiple proteins (60) at complex protein assemblies (such as the KT). For instance, the fuzzy H1-nucleosome complex facilitates the evasion of the disordered chaperone ProTα for the dissociation of H1 from the nucleosome (61). The ultra-high affinity disordered complex formed between H1 and ProTα provides a mechanism for competitive substitution (43).

Scm3 is a compact IDP with several long-range intramolecular interactions that are wrecked by the Cse4 N-terminus to give it a relatively more open and accessible conformation. Additionally, the N-terminus suppresses the conformational exchange (∼R_ex_) of Scm3 thereby stabilizing it. This would enable Scm3 to effectively interact with N-terminus at multiple sites. The relatively accessible conformation of Scm3 in presence of the Cse4 N-terminus may allow interactions with other proteins such as AO at the centromere. Structure-based autoregulation of centromeric proteins has been reported previously for Mif2 and Cse4 wherein both the proteins are in a more accessible conformation upon binding to their respective partners (21, 62).

It has been demonstrated that the END domain residues (50-60) interact with the core domain of Okp1 asserting its role in kinetochore nucleation (23). In our study, we show that Scm3 also interacts with AO and in fact strengthens the interaction between AO and the N-terminus by 3-folds. All the three proteins make simultaneous contacts with other as seen in the fluorescence-based studies (Fig. 5 and Supplementary Fig. 6) thus interacting in a synergistic manner. No CSP change was observed on the ^15^N N-terminus at the Okp1 binding site in the presence of Scm3 which supports the possibility of Scm3-Cse4 N-terminus-AO complex formation. This observation is significant as 1) It provides a basis for KT stabilization (formation of a tight complex at the base of the KT) 2) It sheds light on an IDP-IDP binding mechanism that facilitates synergistic interactions at multiprotein complexes 3) It provides a case of ‘stabilization without folding’ for IDPs.

Based on our observations, we propose that Scm3 associates with the Cse4 N-terminus at first by a ‘fly-casting’ mechanism (used by many histone chaperones (26)) followed by making specific multisite contacts. The N-terminus then gives Scm3 a more open conformation. Scm3 in a more accessible conformation, can bind to both the N-terminus and AO in a synergistic manner. We infer that these interactions facilitate KT assembly and stability during cell cycle progression. Further, this synergistic interaction at the base of the KT (at the Ame1/Okp1link) may help it to withstand the pulling forces generated from the microtubule depolymerization during cell division. Given the high disorder content predicted for KT proteins (Table 2, 14/19 KT proteins), such an interaction mechanism may be common.

## Acknowledgements

We thank Prof. K. Luger (University of Colorado Boulder) for providing the plasmids for WT Cse4, H3, Scm3 and Prof. Westermann (University of Duisburg-Essen) for the plasmid of Ame1/Okp1. We want to sincerely thank Prof. Alex Holehouse for assisting us with computing the kappa Z-scores of the protein sequences. We acknowledge the HF-NMR facility funded by the Research Infrastructure Facility Committee, Industrial Research and Consultancy Centre, Indian Institute of Technology (IIT) Bombay for NMR time; We are also grateful to DBT (BT/PR21656/BRB/10/1562/2016) for project funding and Prof. Ghosh (IITB) for his critical inputs and manuscript proof-reading. SS is grateful for CSIR-UGC, PA for CSIR and AB for UGC Dr. DS-Kothari Postdoctoral fellowship.

---

**Supplementary Table 1:** Experimental parameters for Cse4 N-terminus assignment.

| Experiment | No. of TD points | Spectral width (ppm) | Acquisition time (ms) | No. scans | Total acquisition time |
| --- | --- | --- | --- | --- | --- |

|  | t<sub>3</sub> | t<sub>3</sub> | t<sub>1</sub> | ω<sub>3</sub> | ω<sub>2</sub> | ω<sub>1</sub> | t<sub>3max</sub> | t<sub>3max</sub> | t<sub>1max</sub> |  |  |
| --- | --- | --- | --- | --- | --- | --- | --- | --- | --- | --- | --- |
| HNCO | 2048 | 40 | 112 | 12 | 24 | 20 | 113.6 | 11 | 14.8 | 16 | 24 h |
| HN(CA)CO | 2048 | 40 | 112 | 12 | 24 | 20 | 113.6 | 11 | 14.8 | 16 | 24 h |
| HNCACB | 2048 | 56 | 128 | 12 | 24 | 64 | 113.6 | 15.3 | 5.3 | 16 | 1 day 14 h |
| HN(CO)CACB | 2048 | 56 | 128 | 12 | 24 | 66 | 113.6 | 15.3 | 5.1 | 16 | 1 day 20 h |
| HNN | 1024 | 60 | 80 | 12 | 24 | 24 | 71 | 20 | 27 | 32 | 20 h |
| CC(CO)NH | 2048 | 56 | 112 | 12 | 24 | 66 | 113.6 | 15.3 | 4.5 | 16 | 1 day 14 h |
| HCC(CO)NH | 2048 | 56 | 112 | 12 | 24 | 12 | 113.6 | 15.3 | 6.2 | 16 | 1 day 14 h |

