## Supplementary material for "Intrinsic disorder in CENP-A^Cse4^ tail and its chaperone facilitates synergistic association for kinetochore stabilization": Supplementary Figures_bioarchive.pdf

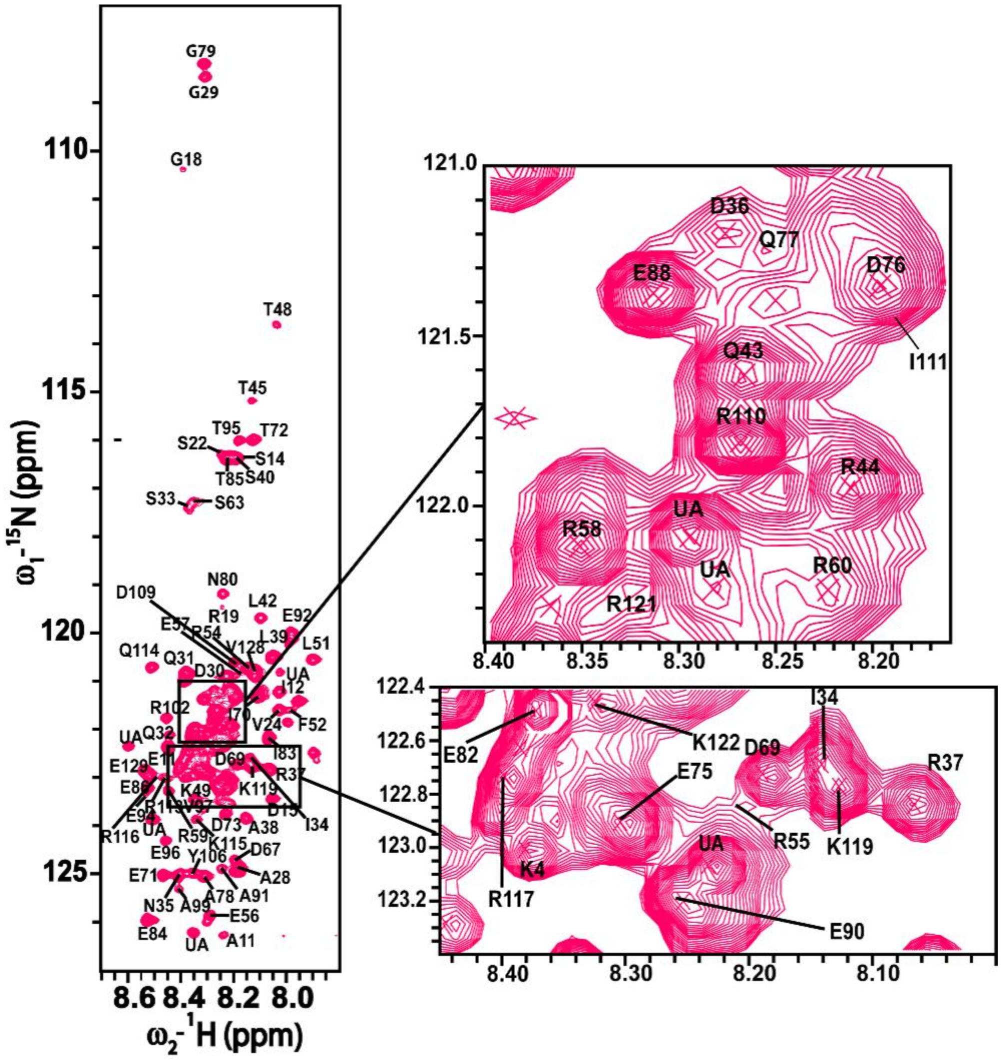


**Supplementary Fig. 1: Resonance assignment of Cse4 N-terminus:** Assigned $^{15}N$-$^{1}H$ resonances in the HSQC spectrum of Cse4 N-terminus at 298 K. The cross peaks are labeled with single letter amino acid code and their corresponding number in the sequence. Backbone assignments for 103/129 peaks could be obtained.

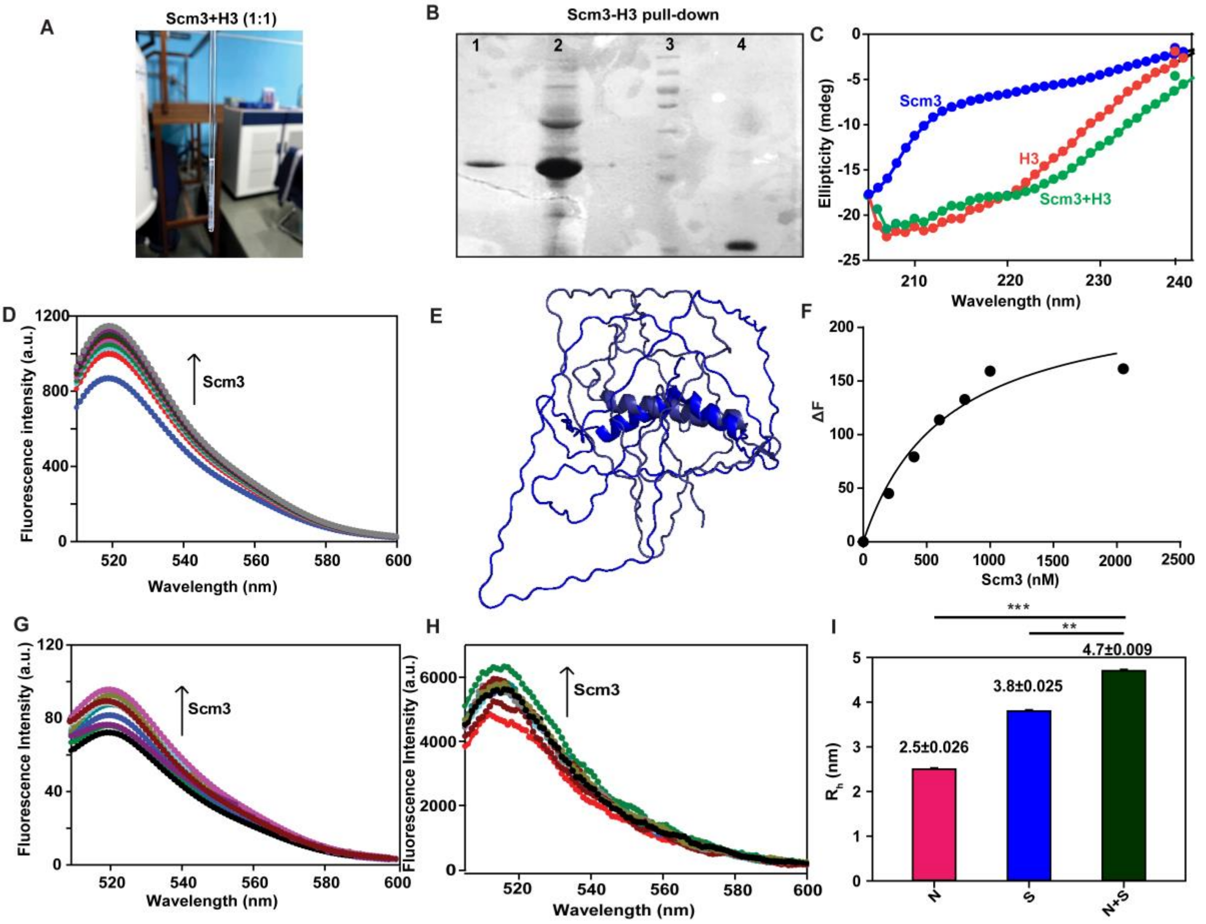


**Supplementary Fig 2: Cse4 N-terminus interacts with Scm3**: A) Equimolar mixture of Scm3 and H3 at 200 µM shows a white precipitate B) 12% SDS-PAGE showing *in vitro* pull down product of Scm3 with H3 (Lane 1), Lane 2 indicates Scm3 in lysate bound to resin used as bait, Lane 4 has the protein molecular weight marker and Lane 5 has pure H3 for reference C) Far-UV CD profiles of H3 (red), Scm3 (royal blue) and complex (green) D) Fluorescence spectra of 100nM FITC-N-terminus in absence and presence of increasing concentrations of Scm3 (100-5000 nM). The upward arrow indicates increasing concentrations of Scm3 E) NMR ensemble of Scm3 as generated by CS23D2.0 E) Scm3 ensemble generated by CS23D2.0 using $^{1}H$, $^{15}N$ and $^{13}C$ NMR chemical shifts of Scm3 F) Binding isotherm of Cse4 in presence of increasing concentrations of Scm3 for $k_d$ determination G) Fluorescence spectra of 100nM FITC-H3 in absence and presence of increasing concentrations of Scm3 (100-5000 nM). The upward arrow indicates increasing concentrations of Scm3 H) Fluorescence spectra of 100nM FITC-α-synuclein in absence and presence of increasing concentrations of Scm3 (100-20000 nM). The upward arrow indicates increasing concentrations of Scm3 I) Bar plot representing the $R_H$ values of N-terminus (magenta), Scm3 (royal blue) and 1:1 complex (bottle green) calculated from the Diffusion constant (D) obtained from DOSY-NMR.

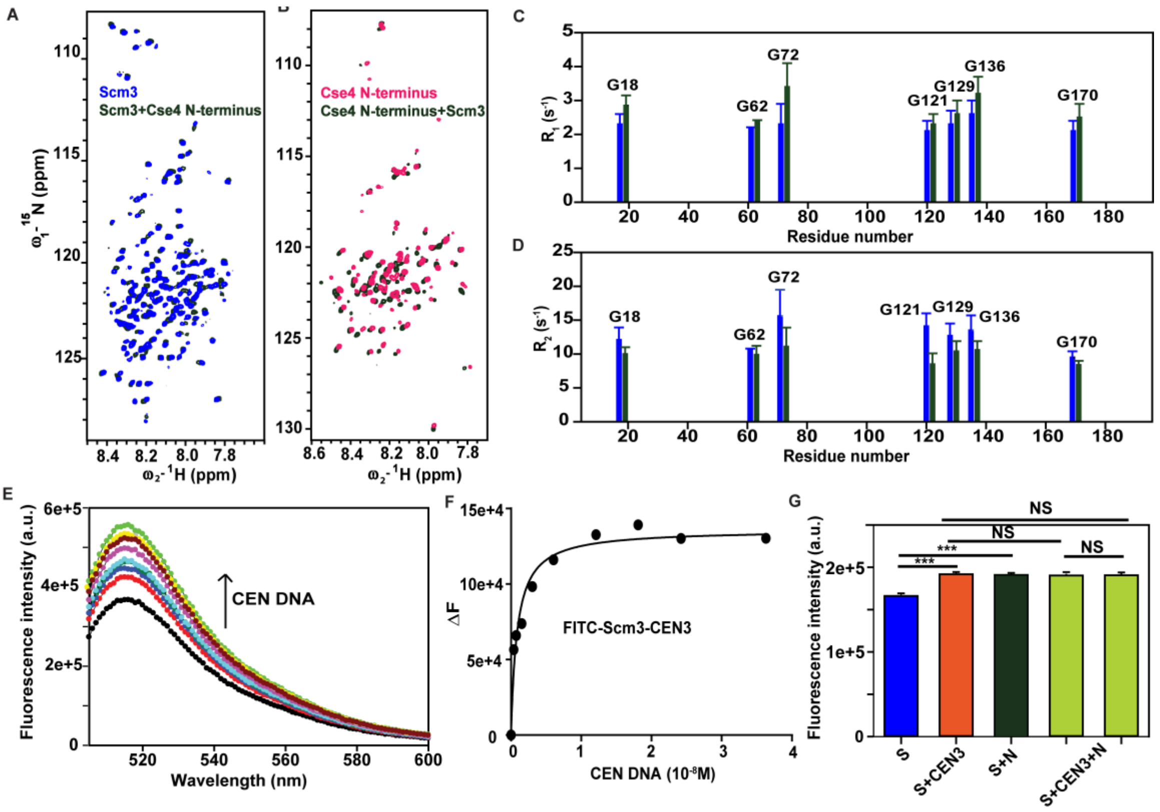


**Supplementary Fig 3: Scm3 specifically interacts with N-terminus:** A) $^{15}$N-$^{1}$H HSQC spectra of $^{15}$N Scm3 (royal blue) and $^{15}$N Scm3 in presence of N-terminus (bottle green) B) $^{15}$N-$^{1}$H HSQC spectra of $^{15}$N N-terminus (magenta) and $^{15}$N N-terminus in presence of Scm3 (bottle green) C) $R_1$ of Glycines of $^{15}$N Scm3 (royal blue) and $^{15}$N Scm3 in presence of N-terminus (bottle green) D) $R_2$ of Glycines of $^{15}$N Scm3 (royal blue) and $^{15}$N Scm3 in presence of N-terminus (bottle green) E) Fluorescence spectra of 100nM FITC-Scm3 in absence and presence of increasing concentrations of CEN3-DNA (3.8-363 nM).The upward arrow indicates increasing concentrations of CEN3-DNA F) Binding isotherm of Scm3 in presence of increasing concentrations of CEN3 for $k_d$ determination G) Competition assay of Scm3-CEN DNA and N-terminus: Blue bar represents the fluorescence intensity maxima of 100 nM FITC-Scm3 denoted as S (royal blue), 100 nM FITC-Scm3+ 30 nM CEN3 complex denoted as S+CEN3 (orange), 100 nM FITC- Scm3+1000 nM N-terminus complex denoted as S+N (bottle green), 100 nM FITC-Scm3+ 30 nM CEN3 complex in presence of 200 and 400 nM of N-terminus respectively denoted as S+CEN3+N (parrot green).

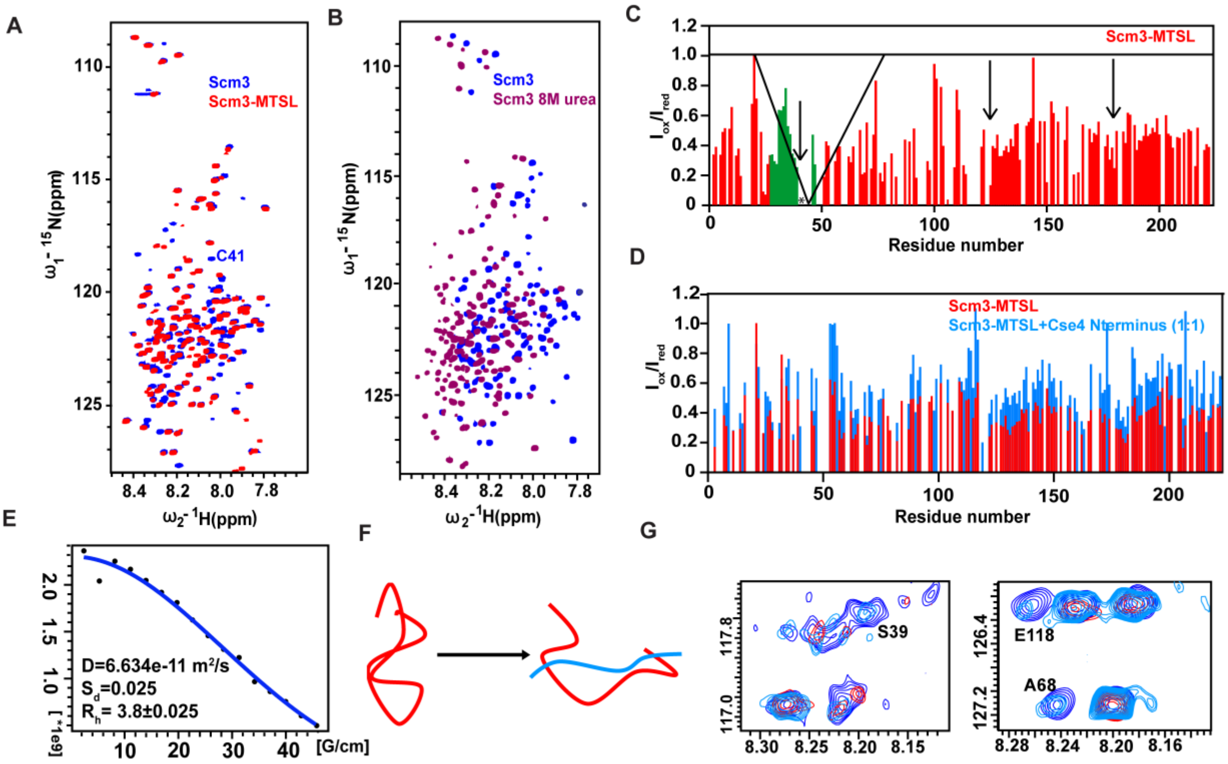


**Supplementary Fig 4: N-terminus disrupts intramolecular contacts of Scm3**: A) $^{15}$N-$^{1}$H HSQC spectra of $^{15}$N Scm3 (royal blue) and $^{15}$N Scm3-MTSL (red). C41 has been labeled in the spectra for reference B) $^{15}$N-$^{1}$H HSQC spectra of $^{15}$N Scm3 (royal blue) and $^{15}$N Scm3 in 8M urea (purple) C) Residue-wise $I_{ox}/I_{red}$ ratio of $^{15}$N Scm3. The region in green is in immediate vicinity of C41-MTSL and shows a visible dip in intensity (downward arrow). * represents C41 and downward arrows represents region of intensity dip D) Fit curve for DOSY-NMR of N-terminus E) Residue-wise $I_{ox}/I_{red}$ ratio of $^{15}$N Scm3 (red) in the presence of N-terminus (blue). Inset: S39 and E118 reappear (blue) in $^{15}$N Scm3-MTSL spectra (red).

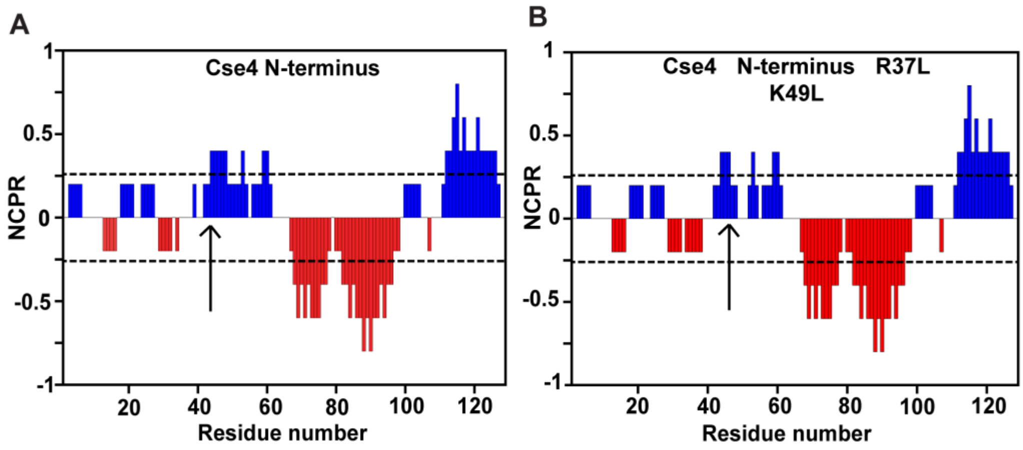


**Supplementary Fig 5: Charge patterning of Cse4 N-terminus in presence of R37 and K49 PTMs:** Local alterations in charge patterning due to R37L K49L has been indicated by the upward arrow. The plot has been generated from CIDER.

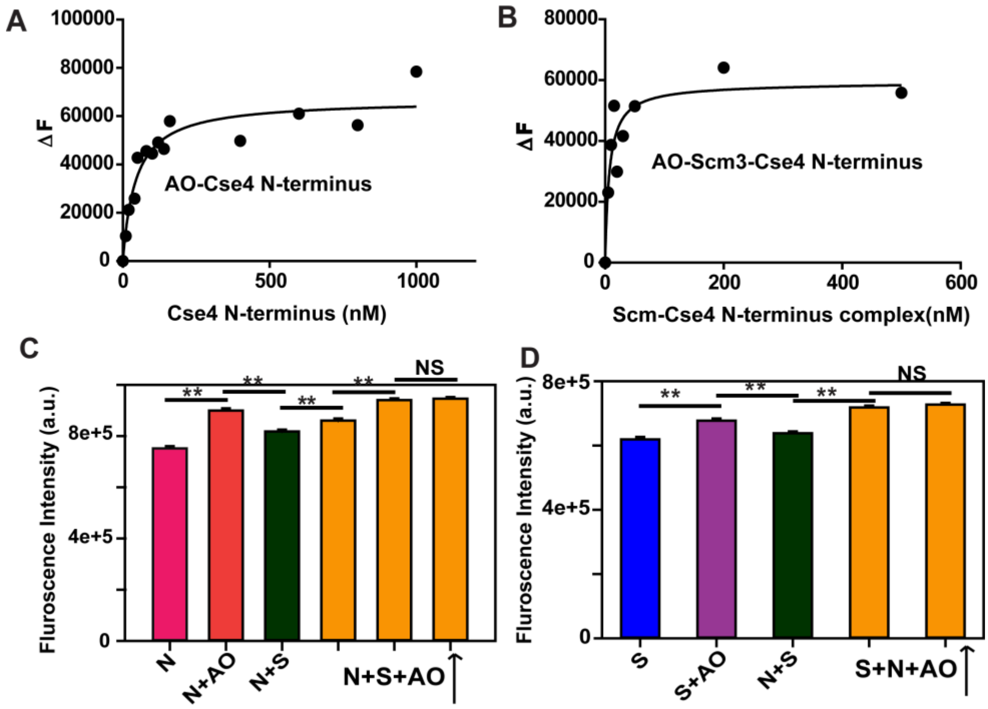


**Supplementary Fig 6: N-terminus-Scm3-AO form a synergistic complex:** A) Binding isotherm of 50nM FITC-AO in absence and presence of increasing concentrations of N-terminus (5-800 nM) for $k_d$ determination B) Binding isotherm of 50nM FITC-AO in absence and presence of increasing concentrations of Scm3-N-terminus complex (5-800 nM) for $k_d$ determination C) Assay of N-terminus-Scm3-AO: Purple bar represents the fluorescence intensity maxima of 50 nM FITC-AO denoted as AO (purple), 50 nM FITC-AO+ 50 nM Scm3 complex denoted as AO+S (dark pink), 50 nM FITC-AO+50 nM N-terminus complex denoted as AO+N (blue), 50 nM FITC-AO+ 50 nM N-terminus complex in presence of increasing concentrations of Scm3 denoted as AO+N+S (orange) D) Assay of N-terminus-Scm3-AO:Pink bar represents the fluorescence intensity maxima of 50 nM FITC-N-terminus denoted as N (pink), 50 nM FITC-N-terminus+ 50 nM AO complex denoted as N+AO (red), 50 nM FITC-N-terminus+50 nM Scm3 complex denoted as N+S (dark green), 50 nM FITC-N-terminus+ 50 nM Scm3 complex in presence of increasing concentrations of Scm3 denoted as N+S+AO (orange).
